# Angiogenic Signaling Counteracts Shear Stress-driven Arterial Patterning

**DOI:** 10.1101/2024.01.23.576920

**Authors:** Dongying Chen, Oleksii S. Rukhlenko, Divyesh Joshi, Martina Rudnicki, Brian G. Coon, Raja Chakraborty, Anna Tuliakova, Elena Ioannou, Kathleen A. Martin, Boris N. Kholodenko, Christiana Ruhrberg, Martin A. Schwartz, Michael Simons

## Abstract

Postnatal vascular morphogenesis is a dynamic process that initially involves angiogenesis to form the capillary bed, followed by its partial remodeling into arteries. Vascular endothelial growth factor A (VEGF hereafter), the primary inducer of angiogenesis, is also implicated in arterial specification in developmental contexts. However, it is unclear why arterial patterning is spatiotemporally segregated from angiogenesis, while postnatal arterial specification in animal models with blocked VEGF signaling remains unstudied. Here, we report that VEGF does not induce arterial fate in the capillaries, instead serving as a physiological brake to attenuate fluid shear stress (FSS)-driven capillary-to-arterial cell fate transition. Mouse models with disrupted VEGF signaling reveal impaired angiogenesis but intact and ectopic arterialization. Mechanistically, mechanosensitive transcription factor Sox17 determines the FSS-arterial program, while VEGF suppresses Sox17 transcription activity at arterial promoters and enhancers sites. Angiogenic signaling, while essential for the initial capillary morphogenesis, inhibits arterial specification, thereby protecting the capillary bed from premature arterialization driven by flow. These findings establish a new paradigm in which precise spatiotemporal coordination of environmental stimuli orchestrates angiogenesis, arterial patterning and capillary maintenance during vascular morphogenesis.

---

Vascular morphogenesis, using postnatal mouse retina as an example, involves several complementary processes: expansion of the capillary bed by vessel sprouting, venous differentiation, and remodeling of the capillary bed into arteries in a process termed arterial patterning^1–3^. Vascular endothelial growth factor VEGF-A (VEGF hereafter) is a key driver of vascular growth^2–6^. VEGF’s role as the primary driver of sprouting angiogenesis has been clearly established, including its role in inducing filopodia-studded endothelial tip cells that lead new vessel sprouts^6,7^, and its crosstalk with DLL4-driven notch signaling^8,9^. Crosstalk of VEGF and Notch signaling are also implicated in promoting arterial specification in specific developmental contexts^10–13^. For example, studies of dorsal aorta formation in zebrafish and mouse showed that VEGF induces Dll4, with subsequent Notch activation promoting arterial specification^10,13,14^. Moreover, it has been suggested that angiogenesis and arterial specification are functionally linked via Notch signaling^15,16^. Constitutive Notch activation via overexpression of the Notch intracellular domain (NICD) increases arterialization^17^, whereas endothelial-specific knockout of Dll4 or treatment with a gamma secretase inhibitor DAPT that inactivates Notch signaling causes vascular hypersprouting whilst decreasing arterial morphogenesis^8,10,11,15,16,18–20^. Collectively, these findings suggest that VEGF, the primary stimulus for angiogenesis, also promotes arterial fate specification through Notch signaling.

However, contrary to the well characterized angiogenic phenotypes in postnatal VEGF loss-of-function (LOF) mouse lines, the requirement of VEGF-A-VEGFR2 signaling in arterial specification has never been characterized and many questions remain unanswered. In particular, it is unclear why angiogenesis and arterial patterning are spatiotemporally segregated in developing vasculature, as arterialization does not take place at the angiogenic front where VEGF levels are the highest. It is also unclear how VEGF, an endothelial growth factor capable of stimulating cell proliferation, can also promote arterial fate specification, considering that the latter process requires cell cycle arrest^21^. The model of VEGF-induced Notch signaling coupling angiogenesis and arterial fate has not yet been integrated with current knowledge of fluid shear stress (FSS)-induced Notch signaling, which regulates arterial specification and maintains arterial homeostasis^21,22^. If both VEGF and FSS promote arterial fate, the combination of the two inputs additively promote arterialization even in the angiogenic capillary region. Thus, the relative importance and interactions of these two essential environmental guidance cues, VEGF and FSS in arterial patterning needs to be determined.

To address these critical problems, we examined the interactions of VEGF and FSS on arterial specification in vitro and in the developing mouse retina. We report that VEGF signaling serves as a critical physiological brake to attenuate FSS-driven arterial fate acquisition and that loss of VEGF signaling leads to ectopic arterialization of the capillary bed. FSS-driven capillary-to-artery cell fate transition requires cessation of VEGF signaling. Mechanistically, VEGF signaling compromises the transcription activity of Sox17 on arterial promoters and enhancers, thereby deactivating the FSS-induced arterial program. Based on these findings, we propose a novel paradigm on how expansion, patterning and maintenance of the capillary bed are precisely coordinated by the interaction of environmental guidance cues.

## Results

### VEGF counteracts FSS-induced capillary-to-artery endothelial cell fate transition

To test the roles of VEGF and FSS in arterial fate specification during capillary bed remodeling, we used human microvascular blood endothelial cells (HMVBECs, primary capillary endothelial cells) to examine the induction of arterial genes under four experimental conditions: static flow (ST), VEGF in static flow (VEGF), laminar flow (FSS), and a combination of laminar flow and VEGF (FSS+VEGF)(Fig. 1a-c). Analysis of the RNAseq transcriptomic data clustered the arterial and capillary genes separately and revealed a pattern of gene expression inversely regulated by VEGF and FSS (Fig. 1a). FSS, but not VEGF, effectively induced the arterial gene expression profile, including definitive arterial genes Gja4, Gja5, Sox17, Efnb2, Sema3g, Cxcl12, Nrp1 and Jag1, and inhibited the capillary profile (Emcn, Flt4 and Mfsd2a) (Fig. 1a). In contrast, VEGF decreased the FSS-induced arterial genes and maintained the capillary genes (Fig. 1a-c). RNA and protein expression patterns of these genes were validated by RT-PCR and western blotting (Fig. 1b-c), except that changes in capillary genes Tfrc and Ivns1abp did not reach statistical significance (Fig. 1b-c). These results indicate that FSS promotes capillary-to-artery cell fate transition, while VEGF counteracts this process to maintain capillary fate (Fig. 1a, diagram).

**Figure 1.**
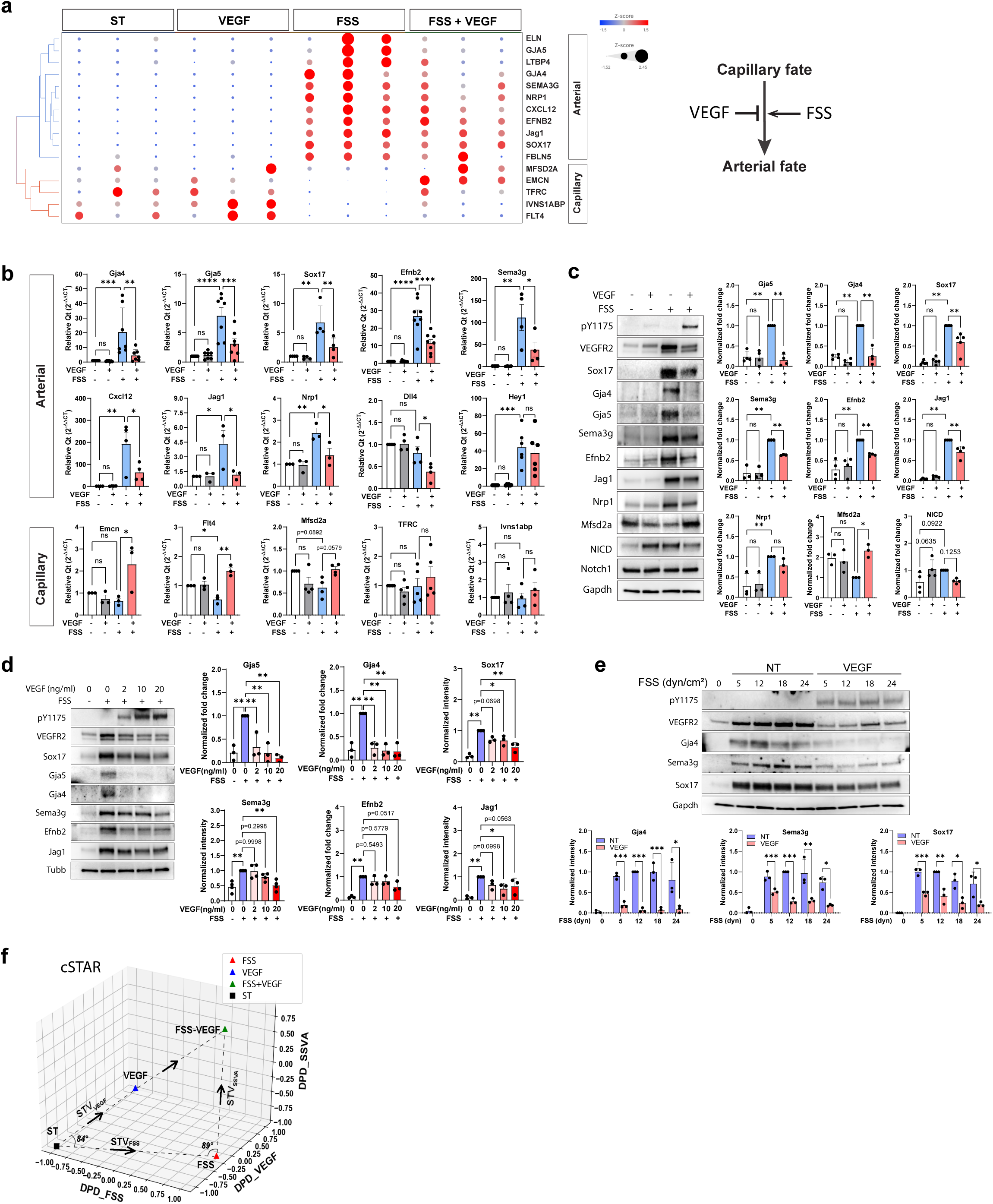
VEGF-A counteracts FSS-driven capillary-to-artery cell state transition. **a.** RNAseq differential gene expression clustering analysis of arterial and capillary genes. HMVBECs were cultured in the conditions of static (ST), VEGF (20ng/ml), FSS (12dyn/cm^2^) and FSS+VEGF for 16 hours. Gene expression matrix is demonstrated by both color intensity and size. **b-c.** Validation of the arterial and capillary genes by quantitative RT-PCR (**b**) and western blot (**c**). **d.** Western blot results of HMVBECs cultured in the conditions of ST, FSS with a gradient of VEGF (0, 2, 10, 20ng/ml). **e.** Western blot results of HMVBECs treated with a gradient of FSS with and without VEGF. **b-e.** One-way ANOVA with multiple comparisons was used for statistics. A p value equal to or less than 0.05 was determined as statistically significant, in extents demonstrated by asterisks * (p≤0.05), ** (p≤0.01), *** (p≤0.001), **** (p≤0.0001). ns: not significant. **f.** cSTAR analysis of RNAseq data. DPD scores of EC states shown in 3D space of the STV_FSS_, STV_VEGF_, and STV_SSVA_. Four EC states correspond to different FSS and VEGF treatments. A change in the EC phenotype is quantified by the DPD score along the corresponding STV. The first cSTAR step uses a machine learning technique called support vector machines (SVM) to exploit the high-dimensional transcriptomics space and construct separating surfaces between distinct EC states. Subsequently, the directions of EC state conversions are determined in the transcriptomic space. These directions are the shortest paths from one EC state to another state, which are specified by the State Transition Vectors (STVs). STVs are unit vectors defined as normal vectors to the hyperplanes that separate EC phenotypic states. After building the STVs, cSTAR calculates quantitative indicators of cell phenotypic states termed Dynamic Phenotype Descriptors (DPD). For EC cells in different states, the DPD scores describe phenotypic features associated with each STV^28,29^. The STV_FSS_ indicates the shortest path from an EC ST state to a FSS state, and the DPD_FSS_ quantifies a normalized FSS-related score of the changes in the transcriptomics patterns. The STV_VEGF_ indicates the direction from an ST state to a new EC state caused by VEGF stimulation at zero FSS, whereas the DPD_VEGF_ scores quantify the extent of the transcriptomics changes related to VEGF stimulation. The angle between two directions of the transcriptomics changes, STV_FSS_ and STV_VEGF_ is 84 degrees. The near orthogonality of these two STVs suggests mutually independent transcriptomic changes between the FSS and VEGF EC states, with any slight deviation from orthogonality indicating a minor overlap in the regulatory pathways induced by FSS or VEGF. To better understand changes in the transcriptomic profile of the FSS EC state brought about by VEGF stimulation, we built the STV_SSVA_ to determine the direction of the EC FSS state change induced by a combined FSS and VEGF stimulation. The resulting STV_SSVA_ appears to be almost orthogonal to both STVs determined above (the angle between the two vectors, STV_SSVA_ and STV_FSS_ is 89 degrees, and the angle between the STV_SSVA_ and the STV_VEGF_ is 87 degrees suggesting that this combination (VEGF+FSS) leads to a new state distinct from both STV_VEGF_ and STV_FSS._ Based on the STV_SSVA_, we next calculated the DPD_SSVA_ that scores the changes in the phenotypic cell features following combined FSS and VEGF treatment. A combined VEGF and FSS treatment only slightly increases the DPD_VEGF_ score of the FSS state and does not change the DPD_FSS_ score.

To test EC subtype generality, we examined human umbilical venous endothelial cells (HUVECs) in the same experimental setting (Fig. S1). The expression patterns of most definitive arterial genes (Gja4, Gja5, Sema3g, Efnb2, Jag1) in HUVECs in response to VEGF, FSS and FSS+VEGF were similar to results in HMVBECs (Fig. S1). There were two exceptions: Sox17 (more induction in FSS+VEGF than FSS) and Nrp1 (no significant induction by VEGF, FSS or FSS+VEGF)(Fig. S1), likely due to the venous and embryonic nature of HUVECs.

To further test the role of VEGF/FSS interaction on arterial specification, we examine the induction of arterial genes in various VEGF and FSS gradients in HMVBECs. VEGF inhibited the FSS-induced arterial program in a dose-dependent manner (Fig. 1d). Among the arterial genes examined, Gja5 and Gja4 were the most sensitive to VEGF inhibition, as the lowest dose (2ng/ml) of VEGF completely blocked their induction by FSS (Fig. 1d). Moreover, induction of arterial genes by FSS gradient (0, 5, 12, 18, 24 dyn/cm^2^) were effectively suppressed by VEGF(Fig. 1e).

To better understand the capillary-to-artery cell state transition, we applied a systems biology approach termed Cell State Transition Assessment and Regulation (cSTAR) analysis^23–25^ to our transcriptomic data (Fig. 1f, detailed description in the legend). cSTAR analysis showed that the four experimental conditions (ST, VEGF, FSS, FSS+VEGF) resulted in four very different metastable EC states (Fig. 1f) in which the FSS+VEGF state was distinct from any of the others. The combination of VEGF and FSS thus induces a distinct program with little overlap with either VEGF alone or FSS alone. Collectively, these findings demonstrate that, during arterial specification, the transition from capillary fate to arterial fate is governed by the interaction between VEGF and FSS: FSS promotes capillary-to-artery transition and VEGF counteracts this process to maintain capillary fate.

### Arterial morphogenesis takes place in the capillary regions with reduced VEGF activity

These data prompted us to re-examine the spatiotemporal relationship between angiogenesis and arterial formation as well as the relationship between VEGF, FSS, and arterial morphogenesis. In the mouse postnatal retina, vascular morphogenesis begins at postnatal day 0(P0)-1 when nascent vessels begin to sprout from the central vein^7,26^, with a rudimentary capillary plexus emerging by P2 (Fig. 2a). At P2-3, when a capillary bed is formed, arteries have appeared in the central part of the capillary plexus, as identified by high expression of the arterial marker Sox17 (Sox17^high^), (Fig. 2a). At P3-4, while sprouting continues at the angiogenic front, arteriovenous patterning transforms the capillary bed into a hierarchical vascular structure with capillaries, arteries and veins (Fig. 2a). By P4-5, the entire vasculature exhibits a stereotypical vascular tree: the arteries and the veins aligned in an alternating pattern in the central area, angiogenic sprouts at the periphery driving vascular expansion towards the avascular zone, and the capillary bed in the middle. (Fig. 2a). Of note, although both angiogenesis and arterial formation progress from the retinal center to the periphery (Fig. 2a, retinal periphery at the top and retinal center at the bottom), extension of the arteries lags, as demonstrated by the un-arterialized gap between the Sox17^high^-labeled arterial tip and the angiogenic front (Fig. 2a). Angiogenesis and arterial patterning are thus spatiotemporally segregated during vascular morphogenesis.

**Figure 2.**
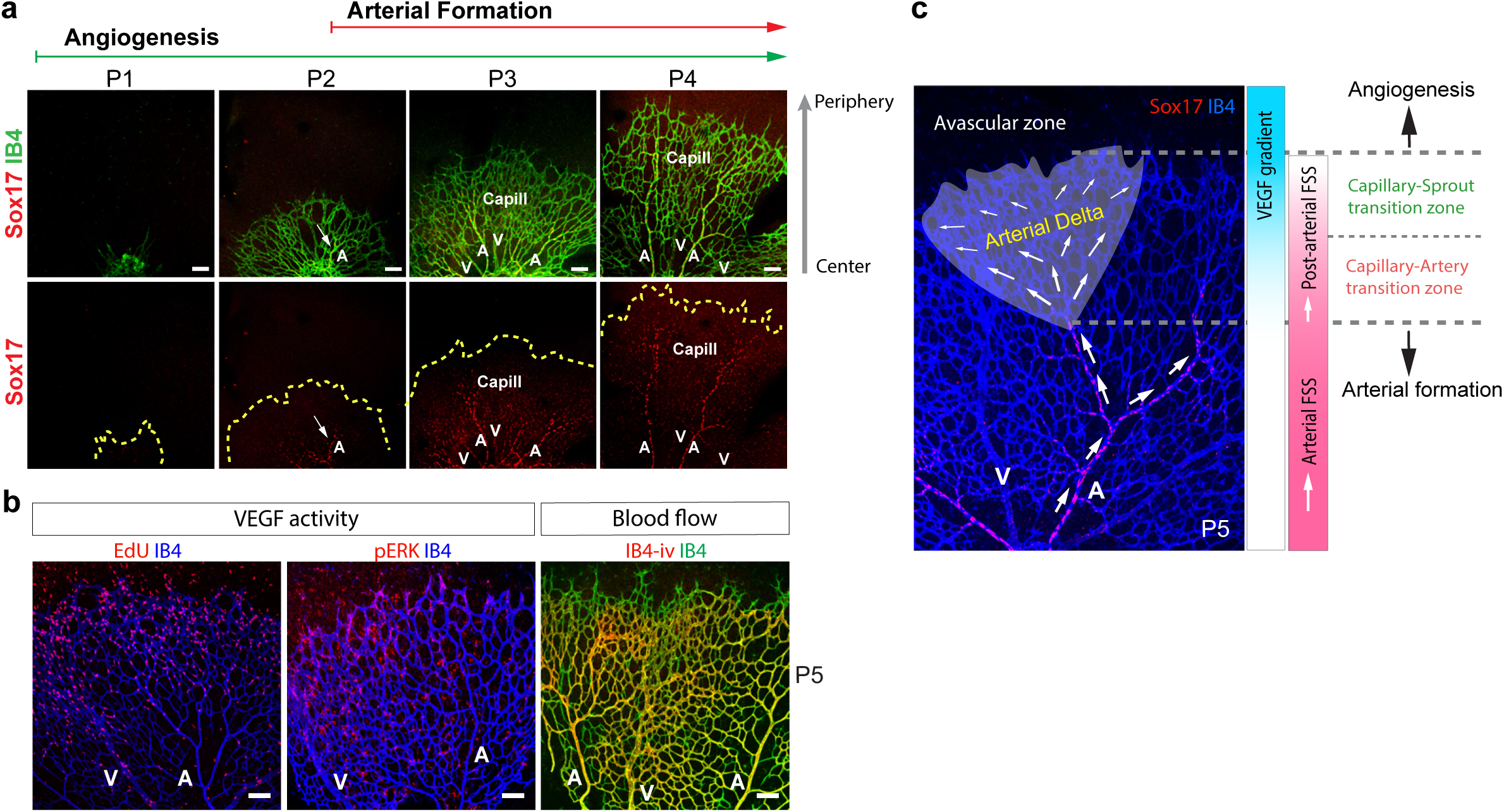
Arterial morphogenesis takes place in the capillary regions with declined VEGF activity. **a.** Formation and remodeling of the capillary bed in mouse postnatal retina. Mouse retinas were dissected at P1-4, followed by immunostaining of Sox17 and IB4. The arrow in P2 indicates emergence of arterial morphogenesis labeled by high level of Sox17 expression. The gray arrow on the right indicates the direction of vascular expansion from the retinal center to the periphery. The yellow dashed lines in the bottom panel represent the dynamic movement of the vascular fronts visualized in the top panel. **b.** VEGF activities and blood flow in P5 retina. i.v.: intravenous (retro-orbital) injection. **c.** A stereotypic vasculature at P5 with immunostaining of Sox17 and IB4. The arrows demonstrate arterial and post-arterial blood flow, from the artery to the un-arterialized artery associated capillary (defined here as Arterial Delta). The right bars demonstrate the gradients of VEGF and FSS. Abbreviations: A: artery; V: vein. Scale bars: 100µm.

In this developing vasculature, EdU incorporation and endothelial phospho-ERK (pERK) signal identified endothelial cells with presumably high VEGF activity, predominantly at the angiogenic front of the P5 retina (Fig. 2b). pERK and EdU-labeled ECs declined in the capillary region undergoing arterialization and almost completely disappeared in arterioles and arteries (Fig. 2b), as previously reported^27^. By contrast, most of the developing vasculature is perfused, visualized by staining with retro-orbitally injected isolectin B4 (IB4-iv), indicating the presence of FSS, except for angiogenic sprouts at the vascular front (Fig. 2b). These observations demonstrate that arterial morphogenesis takes place in the region of the capillary bed under FSS but where VEGF has decreased, and that there is no correlation between high VEGF activity and artery formation.

Intriguingly, the spatial segregation of angiogenesis and arterial formation creates a region between the arterial tips and the angiogenic front (Fig. 2c) that contains organized capillaries but not arteries. We name this region the “*Arterial Delta*” (AΔ), where arterial blood vasculature extends and branches throughout the capillary network in a manner resembling a river delta (Fig. 2c). This region is exposed to both high levels of VEGF and post-arterial levels of FSS, but without arterialization (Fig. 2b-c).

### VEGF signaling is not required for arterial specification during capillary remodeling

The notion that arterial formation occurs in the capillary region where VEGF activity has declined raises the question whether VEGF is in fact required for arterial fate specification. Despite ample studies of angiogenesis, arterial formation and fate specification in VEGF loss-of-function (LOF) mice have never been formally examined. Therefore, we used a comprehensive set of VEGF LOF mouse strains to examine the role of VEGF in arterial formation. Given that arteries derive from capillary remodeling, we defined arterial extension as the ratio of the Sox17^high^ blood vessel (the artery) to the radial outgrowth of the capillary bed labeled by IB4, expressed as a percentage, with 100% representing full extension to the vascular margin. Conversely, the un-arterialized gap from the Sox17^high^ arterial tip to the vascular front was measured and expressed as the percentage of the un-arterialized distance in the radial outgrowth. In addition, mural smooth muscle coverage of the forming artery was examined by SMA antibody staining.

First, we evaluated the effect of inducible global deletion of VEGF, the principal activator of VEGFR2, using the Cag-CreER^TM^ driver to globally delete the *Vegfa* gene (hereafter *Vegfa^iKO^*), as retinal VEGF is mostly produced in the neural bed^4^. Tamoxifen injection to activate the Cre driver was started at P2 when the initial capillary bed is formed but arterial specification is minimal. As expected, early deletion of VEGF significantly reduced vascular outgrowth and capillary density (Fig. 3a). Nevertheless, Sox17 and SMA immunostaining revealed the presence of arteries displaying high Sox17 expression (Sox17^high^) and smooth muscle coverage, both in control and *Vegfa^iKO^* retinas (Fig. 3a). Arterial extension in the control retina was 80.3%±4.8% (mean±SD) with an un-arterialized gap of 19.7%±4.4% (Fig. 1a). In *Vegfa^iKO^*retinas, Sox17^high^-labeled arterial extension was increased (95.1%±5.1%), with ectopic arterialization in the vascular front (indicated by arrow heads) and a strikingly reduced un-arterialized gap (4.9%±5.1%) (Fig. 3a). Of note, 83.3%±14.2% of all individual arteries measured in the *Vegfa^iKO^* retinas reached 100% extension, which was never observed in control retinas (Fig. 1a). Arterial smooth muscle coverage was highly variable in the *Vegfa^iKO^*retinas, with a mildly reduced mean (47.3%±22.8%), compared to 59.5%±9.5% in the control retinas. (Fig. 3a). These observations suggest that, despite mildly impaired smooth muscle coverage, arterial specification in the endothelium remains intact upon global VEGF deletion.

**Figure 3:**
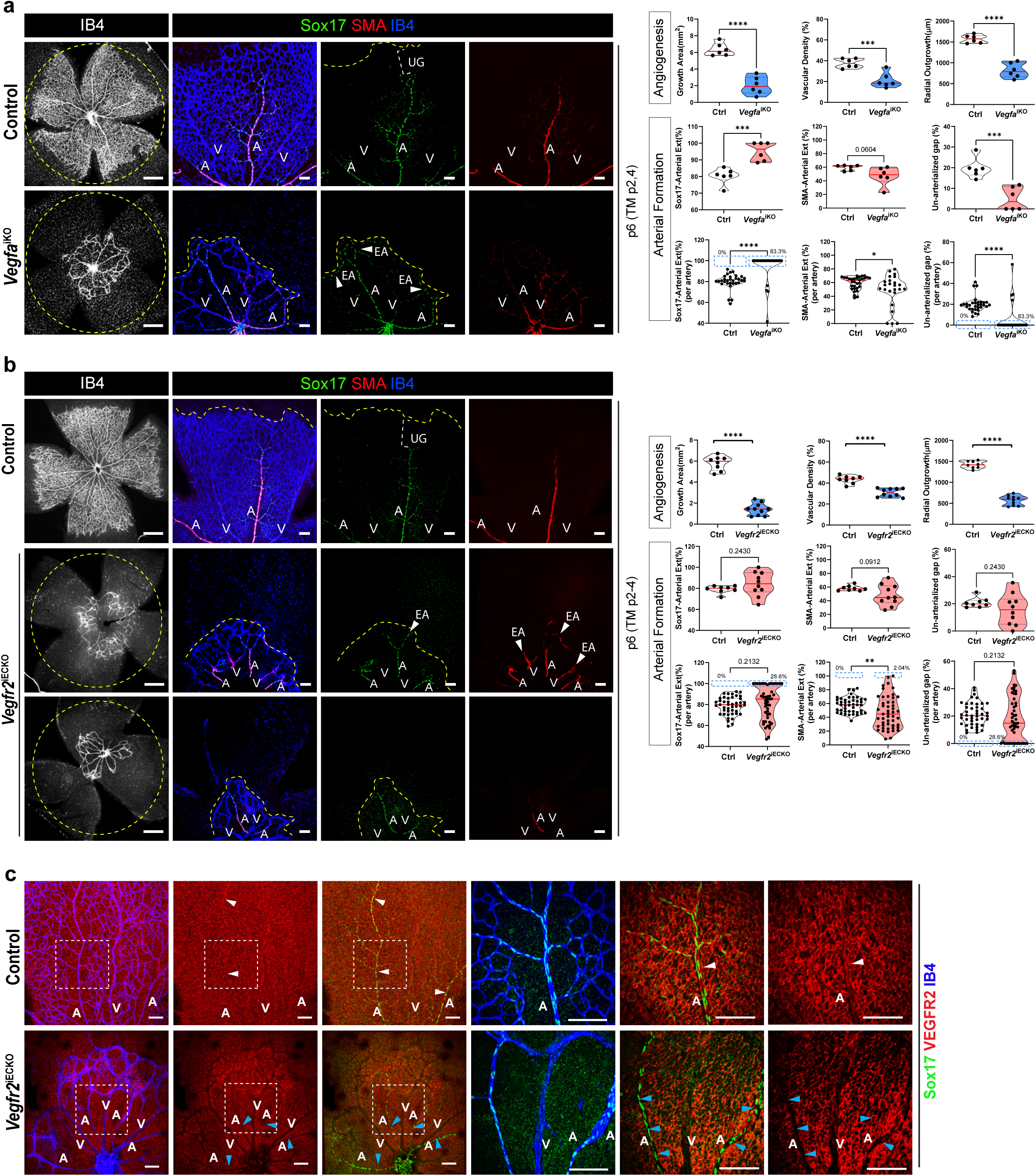
VEGF signaling is not required for arterial specification during capillary remodeling. **a-b**. Mouse neonatal retinas dissected at p6, followed by immunostaining of IB4 and antibodies targeting Sox17 and SMA. Yellow dashed lines indicate the vascular front. White dashed brackets indicate the un-arterialized gap between Sox17-labeled arterial tip and the vascular front. **a**. Control and *Vegfa*^iKO^ retinas dissected at p6. Arrow heads indicate ectopic arterialization (EA). **b**. Control and *Vegfr2^iECKO^* retinas dissected at p6. Blue dashed rectangles in the quantification plots per artery indicate the arteries with full extension and no un-arterialized gap from the artery tip to the vascular front. Abbreviations: A: artery; EA: ectopic arterialization; V: vein. UG: un-arterialized gap. Scale bars: 500 µm (the left columns in a-d); 100µm (the right 3 columns in a-d). Two tailed, unpaired Student t-test was used for statistics. A p value equal to or less than 0.05 was determined as statistically significant, in extents demonstrated by asterisks * (p≤0.05), ** (p≤0.01), *** (p≤0.001), **** (p≤0.0001). Numeric values are reported for p values greater than 0.05. **c.** Control and *VEGFR2^iECKO^*retinas immunostained with IB4 and VEGFR2 to visualize endothelial-specific deletion of VEGFR2. The areas in the white dashed squares are shown in higher magnification in the images of the next column. White arrow heads indicate Sox17^high^/VEGFR2^+^ ECs in the control mice. Blue arrow heads indicate Sox17^high^/VEGFR2^null^ ECs. Abbreviations: A: artery; V: vein. Scale bars: 300µm(a), 100µm (b and c).

Next, we examined the effect of inducible endothelial deletion of VEGFR2 using the Cdh5CreER^T2^ driver (hereafter *Vegfr2^iECKO^*) with tamoxifen injection similarly initiated at P2 (Fig. 3b). Though mean Sox17^high^-labeled arterial extension in the *Vegf2^iECKO^* retinas (84.2%±10.7%) was comparable to the control (79.2%±3.3%), arterial development was highly variable (Fig. 3b). Individual measurements per artery revealed that 28.6% of the Sox17^high^-labeled arteries reached 100% (full) extension in the *VEGFR2^iECKO^*retinas, with ectopic arterialization at the vascular front, which was not observed in the control retinas (Fig. 3b). Mural muscularization also exhibited high variability in the *VEGFR2^iECKO^* retinas, with an average (48.1%±14.6%) mildly reduced compared with the control (58.1%±3.6%) (Fig. 3b). In addition, ectopic muscularization was detected at the vascular front and veins in some *Vegfr2^iECKO^* retinas. Overall, arterial specification is intact in the *VEGFR2^iECKO^* retinas, in line with *Vegfa^iKO^* retinas. Immunostaining with anti-VEGFR2 antibody revealed specific and nearly complete deletion of VEGFR2 in the endothelium but not the neural bed in the *VEGFR2^iECKO^* retinas, creating an appearance of “empty grooves” that highlights the VEGFR2^null^/Sox17^high^ ECs in a VEGFR2+ background (Fig. 3c), direct evidence of VEGFR2-independent arterial specification.

Taken together, these observations suggest that VEGF/VEGFR2 signaling is not required for the acquisition of arterial fate by the capillary endothelium.

### Blocking VEGF signaling leads to ectopic arterialization in the Arterial Delta

Although arterial specification remains intact in the *Vegfa^iKO^* and *VEGFR2^iECKO^*retinas, severe vascular rarefaction in these mice may directly or indirectly affect global and local hemodynamic force, endothelial and mural cell physiology, and subsequent arterial maturation. Also, global loss of VEGF in the *Vegfa^iKO^*retinas potentially impairs VEGF-VEGFR1 signaling in pericytes^28,29^. Therefore, we used alternative approaches to block endothelial VEGF signaling without deleting the ligand and receptor, thereby leaving VEGF and VEGFR2 intact and available for other biological activities.

First, we tested DC101, a well-studied neutralizing antibody that specifically blocks the binding of murine VEGF to VEGFR2^30,31^. DC101 was injected intra-peritoneally into neonatal mice at P1 and P3. DC101 treatment significantly impaired vascular growth and density, confirming effective blockade of VEGFR2 signaling, while substantially preserving the capillary bed (Fig. 4a). Notably, immunostaining of Sox17 and SMA revealed extensive arterialization in the developing vasculature (Fig. 4a). All arteries in the DC101-treated retina spanned from the retinal center to the vascular front, reaching 100% arterial extension, with complete disappearance of the un-arterialized gap (Fig. 4a). Ectopic arterialization, marked by both Sox17^high^ and SMA signals, was found in the Arterial Delta region, leaving only the capillary region between the main arteries and veins un-arterialized across the vasculature (Fig. 4a).

**Figure 4:**
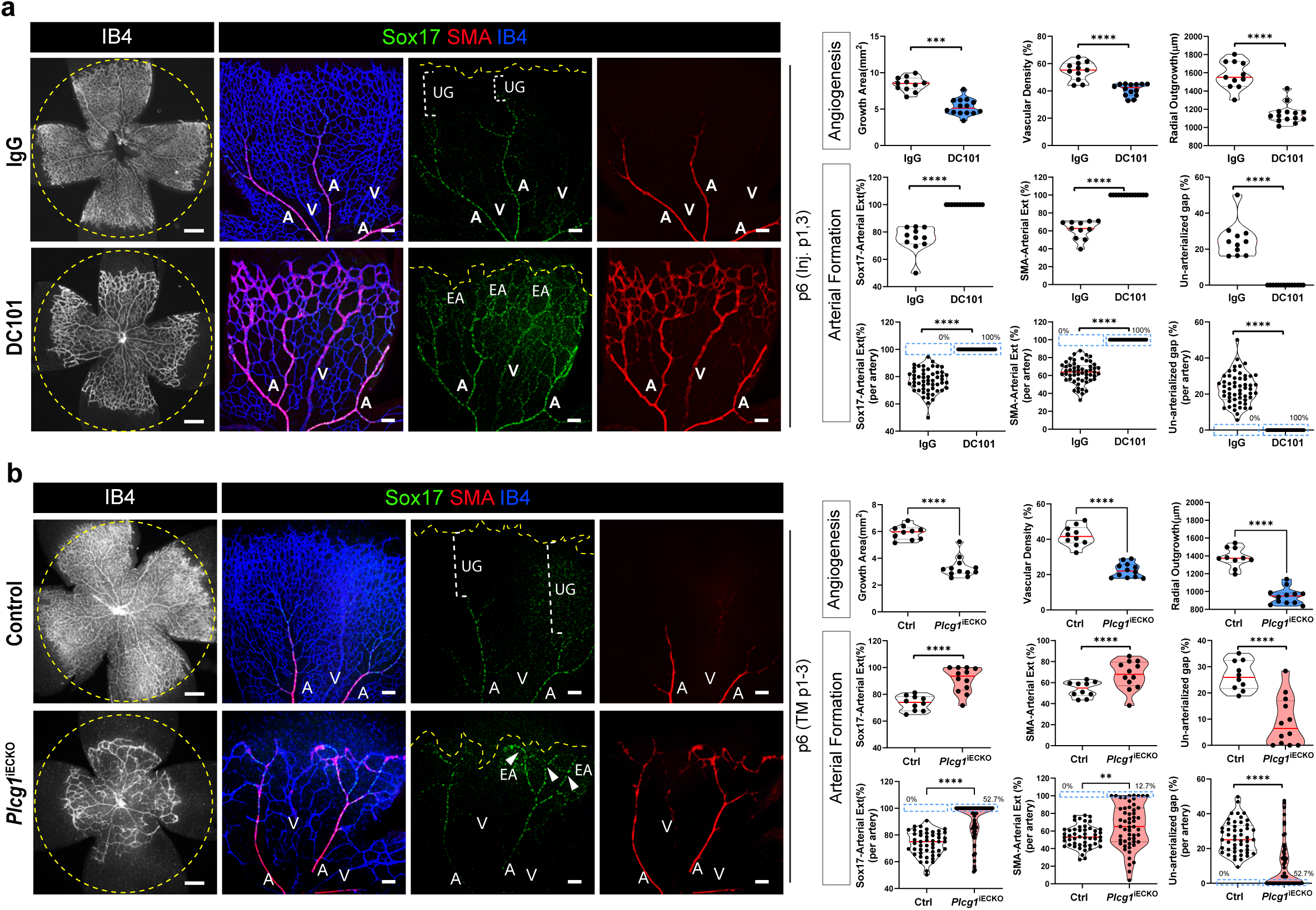
Blocking VEGF signaling leads to ectopic arterialization in the Arterial Delta region. Mouse neonatal retinas dissected at p6, followed by immunostaining of IB4 and antibodies targeting Sox17 and SMA. Yellow dashed lines indicate the vascular front. White dashed brackets indicate the un-arterialized gap between Sox17-labeled arterial tip and the vascular front. **a.** Retinas from pups treated with two doses of IgG or DC101 antibody injection at p1 (400µg) and p3(200µg). **b**. Control and *Plcg1^iECKO^* retinas (tamoxifen induction of gene deletion at p1-3). Blue dashed rectangles in the quantification plots per artery indicate the arteries with full extension and no un-arterialized gap from the artery tip to the vascular front. Arrow heads indicate ectopic arterialization (EA). Abbreviations: A: artery; EA: ectopic arterialization; V: vein. UG: un-arterialized gap. Scale bars: 500 µm (the left columns in a-d); 100µm (the right 3 columns in a-d). Two tailed, unpaired Student t-test was used for statistics. A p value equal to or less than 0.05 was determined as statistically significant, in extents demonstrated by asterisks * (p≤0.05), ** (p≤0.01), *** (p≤0.001), **** (p≤0.0001). Numeric values are reported for p values greater than 0.05.

Next, we tested the effect of endothelial-specific deletion of *Plcg1*. While PLCγ1 mediates multiple signaling pathways, it is the primary effector of VEGFR2 signaling and plays only subordinate roles in other growth factors signaling^32,33^. Endothelial deletion of PLCγ1 (hereafter *Plcg1^iECKO^*) at P1-3 significantly impaired angiogenesis at P6, but with better preservation of the vascular bed(Fig. 4b) compared with the *Vegfa^iKO^*and *VEGFR2^iECKO^* retinas (Fig. 3). Arterial extension was remarkably increased in the *Plcg1^iECKO^* retinas, with 52.7% of Sox17^high^-labeled arteries reaching 100% extension (Fig. 4b). Ectopic arterialization marked by both Sox17^high^ and SMA signals was evident in the Arterial Delta region in *Plcg1^iECKO^*retinas (Fig. 4b). These arterial phenotypes are also in line with the DC101-treated retinas, indicating increased arterial specification with mature mural muscularization.

Due to the lack of a workable staining antibody, deletion of endothelial PLC*γ*1 in P6 pups were analyzed using isolated brain ECs in order to acquire enough cells for qPCR. The results demonstrate 70% reduction of *plcg1* mRNA as detected by the primers targeting exons2-3 and comparable levels of *plcg1* mRNA by the primers targeting exons 24-26, which confirms effective gene deletion of plcg1 despite the presence of a truncated plcg1 mRNA (Fig. S2).

In summary, DC101 treatment and endothelial deletion of PLCγ1 effectively block VEGFR2 signaling, which leads to reduced angiogenesis but intact arterial formation with ectopic arterialization the Arterial Delta region.

### Fluid shear stress (FSS)-induced arterial specification is independent of VEGFR2 signaling

Since arterial specification is intact in the VEGF LOF animal models, we investigated whether FSS is capable of driving arterial fate in the absence of VEGF signaling. FSS requires the presence of blood flow and, hence, vascular lumenization. We retro-orbitally injected IB4-Alexa488 (IB4-iv) to visualize vascular patency by indicating the presence of blood flow (Fig. 5a). Dissected retinas were counterstained with IB4-Alexa647 to visualize the entire vasculature, including closed vessels (Fig. 5a). In control retinas, while the majority of vessels were double-stained, IB4-Alexa488 was absent from un-lumenized sprouts at the vascular front and some regressed vessels around the arteries (Fig. 5a), indicating a lack of blood flow in these vessels. Importantly, the DC101-treated, *Vegfr2*^iECKO^ and the *Plcg1*^iECKO^ vessels were well double-stained (Fig. 2a), suggesting that there was effective blood flow through these vessels and that endothelial cells with defective VEGFR2 signaling were exposed to the hemodynamic force (FSS).

**Figure 5:**
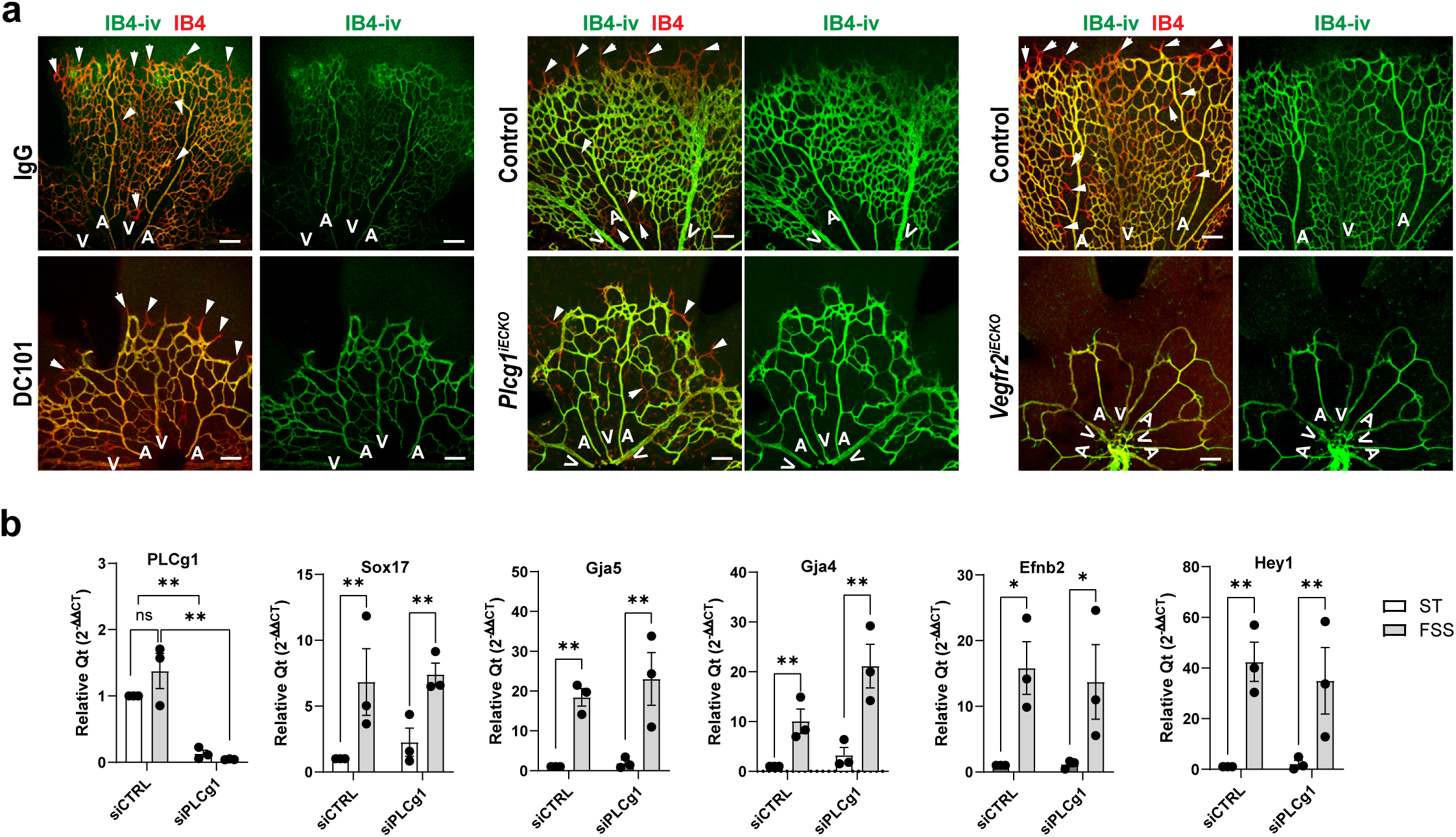
Fluid shear stress-induced arterial program is independent of VEGFR2 signaling. **a**. Mouse postnatal retinas were dissected at p6. Before dissection, IB4 (green, IB4-iv) was retro-orbitally injected to perfuse the retina vasculatures of Control, DC101-treated, *Vegfr2^iECKO^* and *Plcg1^iECKO^*mice. Dissected retinas were stained with IB4 (red) to visualize all perfused and non-perfused blood vessels. Perfused vessels are double-stained (IB4-green/red), while vessels not perfused were single-stained (IB4-red only, indicated by white arrow heads). Abbreviations: i.v.: intravenous (retro-orbital) injection. A: artery; V: vein. Scale bars: 100µm. **b**. HUVECs with siPLCg1 treatments were subjected to static (ST) or FSS (12dyn/cm^2^) for 16 hours to induce arterial differentiation. Gene expression was examined by quantitative RT-PCR followed by quantification relative to ST-siCTRL. Two-way ANOVA with multiple comparisons was used for statistics. A p value equal to or less than 0.05 was determined as statistically significant, in extents demonstrated by asterisks * (p≤0.05), ** (p≤0.01). ns: not significant. Error bars indicate standard error of the mean.

We next tested whether blocking VEGFR2 signaling affects FSS-induced arterial specification in ECs in vitro. HUVECs treated with siPLCγ1 responded to arterial level of laminar FSS (12 dynes/cm^2^) with effective induction of arterial genes, including Sox17, Gja4, Gja5, Efnb2 and the Notch target Hey1(Fig. 5b), indicating an intact FSS-induced arterial program upon blocked VEGF signaling. Taken together, these in vivo and in vitro findings indicate that FSS-induced arterial specification is independent of VEGFR2 signaling, consistent with intact arterial specification in VEGF LOF mouse models.

### Blocking VEGFR2 signaling promotes capillary-to-artery transition in the single cell transcriptome

To gain mechanistic insights into VEGF LOF arterial phenotypes, we performed single-cell RNAseq analysis of endothelial cells (ECs) from control and *Plcg1^iECKO^* mice retinas. Among 30 cell clusters identified in UMAP (uniform manifold approximation and projection) plots, 1505 cells in cluster 12 with endothelial identity (*Pecam1+*, *Cdh5+; Cldn5+*) were selected for further analysis (Fig. S4). Published markers of endothelial phenotypes (Table S1) were used to annotate endothelial zonation along the arterial-capillary-venous axis based on gene expression profiles (Fig. S5). The following 10 EC subclusters were identified (0-9) (Fig. S5a and Fig. 6a): a sprouting cluster (cluster 6, *Esm1^high^;Kcne3^high^*), an arterial cluster (cluster 9, *Gja4 ^high^; Gja5^high^; Efnb2^high^; Sema3g^high^; Gkn3^high^; Bmx^high^; Sox17^high^; Cxcl12^high^; Unc5b^high^*, most cells in G1 phase), capillary/venous clusters (clusters 0, 2, 3, 4 and 7, *Aplnr^high^; Mfsd2a^high^; Ivns1abp^high^; Flt4^high^; Tfrc ^high^; slc38a5 ^high^; Nr2f2 ^high^; Nrp2 ^high^; Ephb4 ^high^*), an arterial predecessor cluster (cluster 8, with both arterial and capillary profiles) and a proliferative cluster (cluster 0, with capillary/venous profile, all cells in G2M/S phases) (Fig. S5b-f). In addition, the ECs in cluster 5 display weak endothelial identity with low expression of endothelial markers (Cdh5, Cldn5, Pecam1, Flt1, Kdr and Erg), likely ECs from regressing vessels (Fig. S5e). The identity of Cluster 1 is unclear, as these ECs express low levels of capillary/venous markers and some endothelial markers (Cdh5, Cldn5 and Kdr) (Fig. S5d-e).

**Figure 6:**
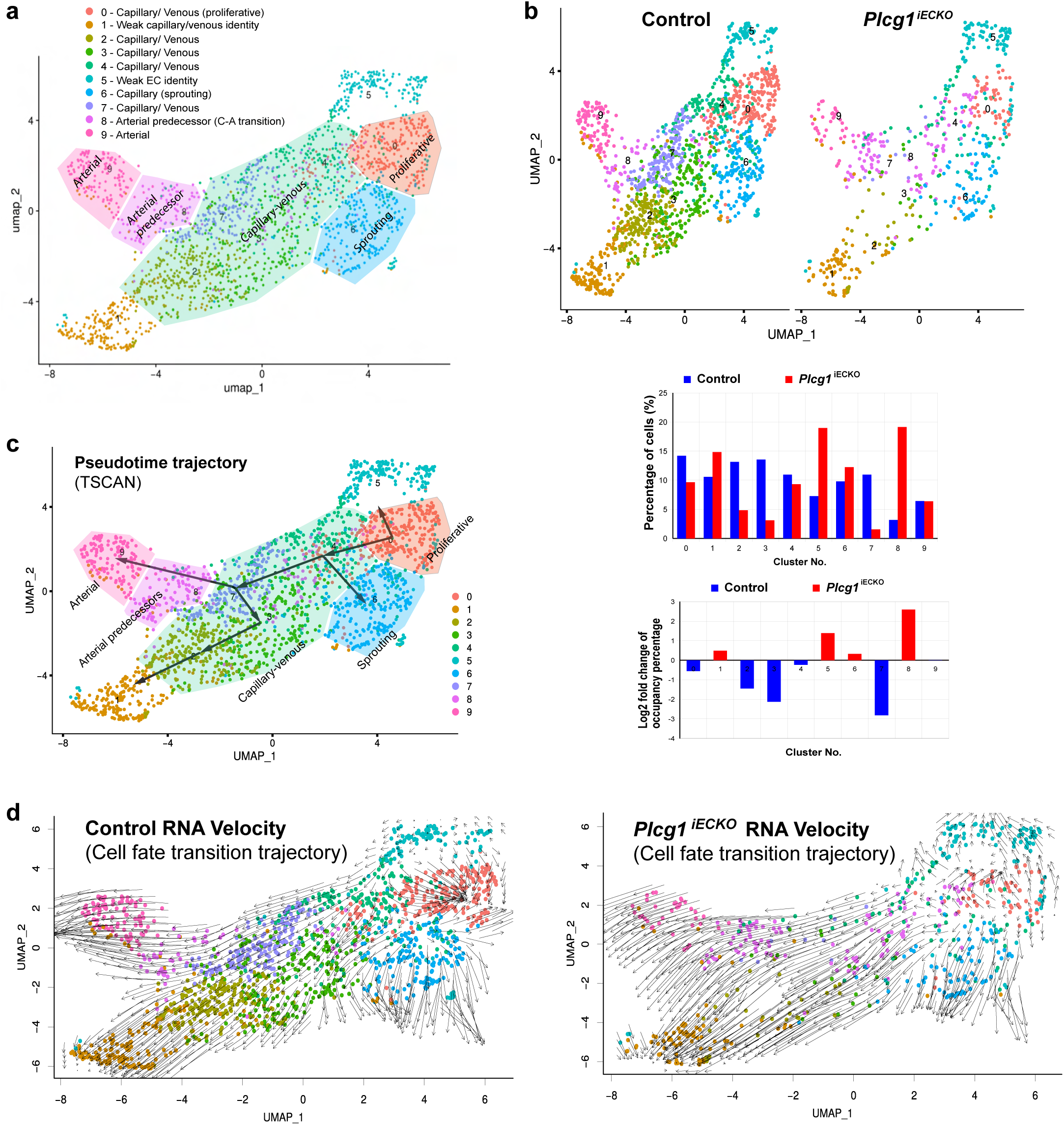
Blocking VEGFR2 signaling promotes capillary-to-artery transition in scRNAseq. **a**. Arteriovenous zonation in combined control and *Plcg1^iECKO^* retinal ECs. **b**. Distributions of control and *Plcg1^iECKO^* retinal ECs in clusters 0-9. Quantification plots demonstrating percentage and log2 fold change of the cells in clusters 0-9 in control and *Plcg1^iECKO^*. **c**. Pseudotime trajectory analysis by TSCAN. **d.** RNA velocity analysis demonstrating transitioning trajectories of control and *Plcg1^iECKO^* retinal ECs.

In the *Plcg1^iECKO^* transcriptome, the percentages of ECs in the proliferative cluster 0 and capillary clusters 2, 3 and 7 were substantially reduced (Fig. 6b), consistent with defective angiogenesis in *Plcg1^iECKO^* retinas. Notably, the *Plcg1^iECKO^* arterial predecessors in cluster 8 (purple color) increased substantially (19% in *Plcg1^iECKO^* versus 3% in the control), with a pattern of distribution spreading into and mixed with capillary clusters 4 and 7 (Fig. 6b), suggesting increased capillary-to-artery transition in the *Plcg1^iECKO^*.

To test this hypothesis, we performed a trajectory analysis to understand the dynamic transition of differentiation states among EC clusters (Fig. 6c-d). Pseudo-time analysis by TSCAN revealed a major *capillary-to-artery (C-to-A)* trajectory starting from the proliferative cluster 0 to capillary/venous clusters 4 and 7 and ending at the arterial predecessor cluster 8 and the arterial cluster 9 (Fig. 6c). In addition to this main trajectory, there were 3 side trajectories branching from the capillary clusters: proliferative cluster 0 transitioned to cluster 5 with weak EC identity, likely due to vascular regression; capillary/venous cluster 4 transitioned to sprouting cluster 8; and capillary/venous clusters 7 transitioned to clusters 3, 2, 1 (Fig. 6c). Importantly, trajectory analysis using RNA velocity revealed an intact C-to-A differentiation trajectory in the *Plcg1^iECKO^* transcriptomes (Fig. 6d), consistent with the increase of *Plcg1^iECKO^* arterial predecessors in cluster 8 (Fig. 6b) as a result of increased C-to-A transition. In addition to the data in Fig. 2-5, scRNAseq analysis thus independently leads to the conclusion that blocking VEGFR2 signaling promotes C-to-A cell fate transition during capillary remodeling.

### VEGF does not significantly alter FSS-induced cell cycle arrest

FSS-induced cell cycle arrest is a prerequisite for arterial specification^21^. Given the role of VEGF in promoting proliferation, we tested whether it antagonizes the FSS-driven arterial program by altering FSS-induced cell cycle arrest. We first conducted transcriptomic analysis of the major cell cycle regulators in the 4 experimental conditions mentioned above (ST, VEGF, FSS, FSS+VEGF, for 16 hours). Differential gene expression clustering showed a clear demarcation between VEGF- and FSS-regulated cell cycle genes (Fig. S6a). FSS effectively silenced activators for G2/M-phase (Cdk1, Ccnb1, Aurka, Aurkb), S-phase (Ccnd1, cdk4) and S/G2 (Ccna1, cdk2), while upregulating pan-cell cycle inhibitors (cdkn1a, cdkn1b) and G1/S activators (Ccne1, cdk6), demonstrating a clear pattern of G1/S arrest (Fig. S6a). VEGF alone had opposite effects (Fig. S6a). However, except for the downregulation of cdkn1b and cdk6, the FSS+VEGF gene set was similar to FSS alone (Fig. S6a). We validated most of these cell cycle regulators using RT-PCR (Fig. S6b). Consistent with these results, EdU incorporation assay confirmed that FSS effectively silenced proliferation, which was not significantly changed with VEGF+FSS (Fig. S6c).

### Notch inhibition does not affect the inhibitory role of VEGF on arterial specification

Notch signaling is an important feature of the arterial vasculature. Since both VEGF and FSS can induce Notch signaling, we investigated its involvement in VEGF-FSS regulation of arterial specification. To this end, HMVBECs were pretreated with 5µM DAPT for 3h to eliminate Notch activity, confirmed by the near absence of cleaved Notch intracellular domain (NICD) (Fig. S6), then subjected to FSS. Notch inhibition did not affect FSS-induced expression Sox17, Sema3G, and Jag1, indicating that induction of these arterial genes is Notch-independent (Fig. S6). However, induction of Efnb2, Gja5, Gja4 and Dll4 was partially or completely compromised, indicating a requirement of Notch signaling for inducing these arterial genes (Fig. S6). Notably, VEGF still attenuated FSS induction of Gja4, Gja5, Sox17, Sema3g, Efnb2 and Jag1, which was not affected by DAPT (Fig. S6). Therefore, although Notch regulates expression of some of the arterial fate genes, it is not required for VEGF-dependent inhibition of FSS-induced arterial specification.

### Sox17 is a mechanosensitive transcription factor essential for the FSS-induced arterial program

The uniform response of the definitive arterial genes to FSS and VEGF regulation suggests a common transcription control point. To investigate this possibility, we used the bioinformatic tool TRANSFAC to identify transcription factors binding sites in arterial and capillary genes. This analysis identified predicted binding motifs for the transcription factor Sox17 in proximal promoters and/or the introns of all arterial and Notch genes, including Sox17 itself, but not in the capillary genes (Table S3). By contrast, the predicted binding motifs for the Notch transcription regulator Rbpj were not found in arterial genes Sox17 and Gja4 but were present in some capillary genes (Table S3). These findings suggest that Sox17 is a potent transcription regulator of the arterial specification program.

Previous studies showed that endothelial deletion of Sox17 impairs arterial specification in mouse postnatal retina^17,34^, but the role of Sox17 in FSS signaling is unclear. We therefore performed Sox17 loss- and gain-of-function analyses. First, we used CRISPR cas9 to knock out Sox17 in HUVECs to test its requirement for FSS to induce other definitive arterial genes. As predicted, the loss of Sox17 profoundly compromised the FSS-induced arterial program, almost completely blocking induction of Gja5, Gja4, Sema3g, Efnb2 and Nrp1 and reducing Unc5b and Jag1 (Fig. 7a). By contrast, CRISPR cas9-mediated deletion of Sox18, another SoxF factor reported to have a compensatory role to Sox17 in arterial specification in mouse retina^34^, only modestly affected FSS induction of Gja4 and Efnb2 and had no effect on other arterial genes tested here (Fig. 7a). Combined knockout of Sox17 and Sox18 had no further effect on the FSS-arterial program (Fig. 7a). Therefore, Sox17, but not Sox18, mediates the FSS-induced arterial program. Next, we tested whether Sox17 overexpression can bypass FSS to induce the arterial program at static condition, using a lentiviral to doxycycline (Dox)-inducible Sox17 system in HUVECs (Dox-Sox17-HUVECs). Dox-induced Sox17 overexpression effectively activated the arterial program without flow (Fig. 7b), with dose-dependent increases in Gja5, Gja4, Sema3g, Efnb2, Dll4 and Unc5b with increasing Sox17 levels (Fig. 7b). Taken together, these findings identify Sox17 as a key mechanosensitive transcription factor in the FSS-induced arterial program.

**Figure 7:**
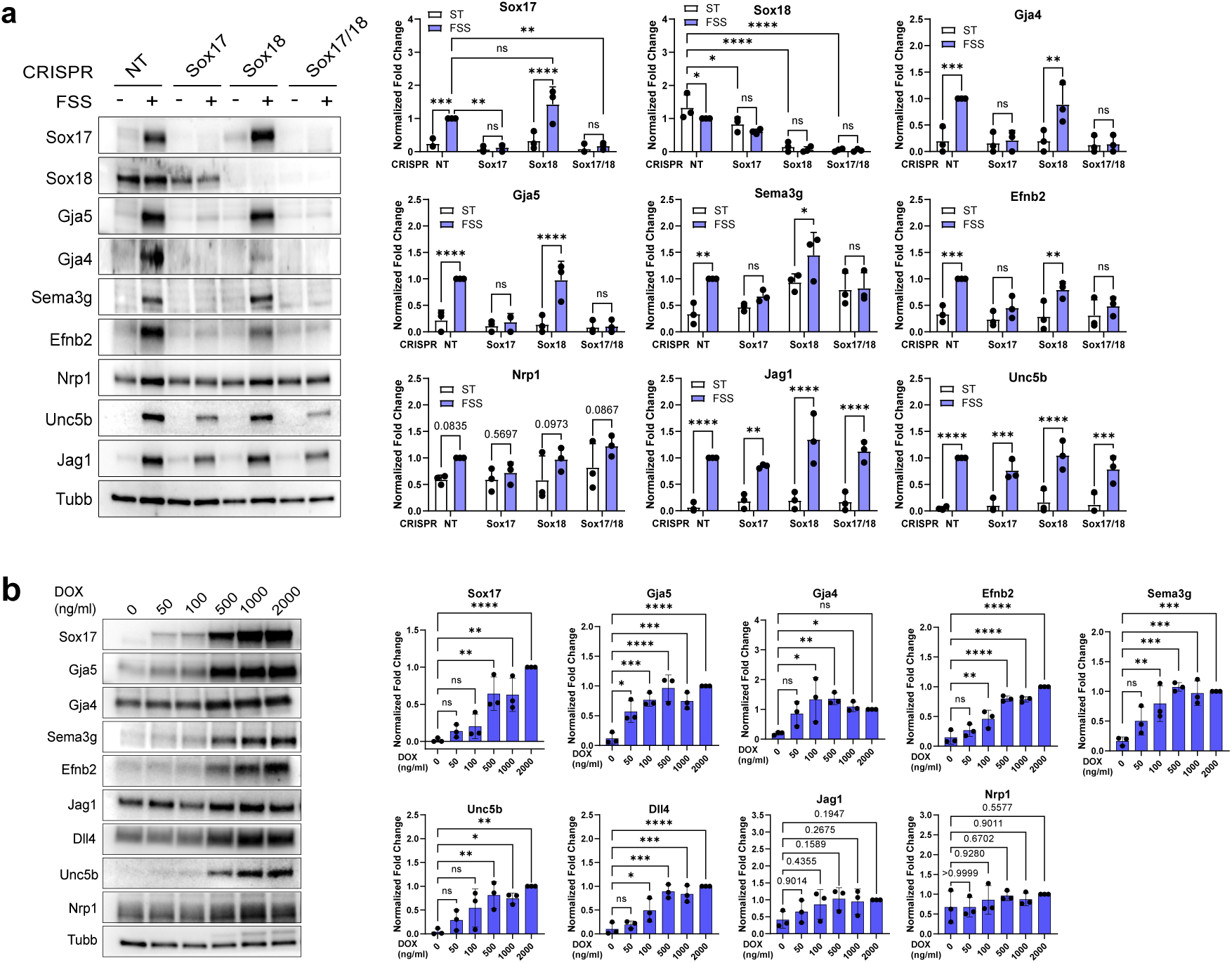
Sox17 is a mechanosensitive transcription factor essential for the FSS-induced arterial program. **a.** HUVECs lentivirally transduced with CRISPR-Cas9-directed sgNT, sgSox17, sgSox18 or combined sgSox17/18 were used to test FSS-induced arterial differentiation. Cell lysates were analyzed by western blot. **b.** HUVECs harboring a doxycycline-inducible Sox17 expression system (pCW57.1-hSox17-myc-DDK) were treated by a gradient of doxycycline (DOX) for 48 hours. Cell lysates were analyzed by western blot. One-way ANOVA with multiple comparisons was used for statistics. A p value equal to or less than 0.05 was determined as statistically significant, in extents demonstrated by asterisks * (p≤0.05), ** (p≤0.01), *** (p≤0.001), **** (p≤0.0001). ns: not significant.

### VEGF antagonizes Sox17 transcription activity on arterial promoters and enhancers

Given the central role played by Sox17 in FSS-dependent arterial fate induction, we next examined whether VEGF antagonizes the FSS program by suppressing Sox17 activity. To this end, we examined the effect of VEGF treatment on arterial gene induction in the Dox-Sox17-HUVEC system discribed above (Fig. 7b). We found that, without lowering the expression level of Sox17, VEGF inhibited Sox17-activated transcription of the definitive arterial genes, including Gja5, Gja4, sema3g, Jag1, Cxcl12 and Efnb2 but did not reduce Sox17 itself (Fig. 8a). These effects depended on VEGF dose (Fig. 8b), similar to the dose-dependent effect of VEGF on inhibiting the FSS-arterial program (Fig. 1d). At the same time, VEGF enhanced expression of Unc5b and Dll4, two known VEGF-Notch targets enriched in both arterial and sprouting ECs^8,35^ (Fig. 8a), providing a positive control.

**Figure 8:**
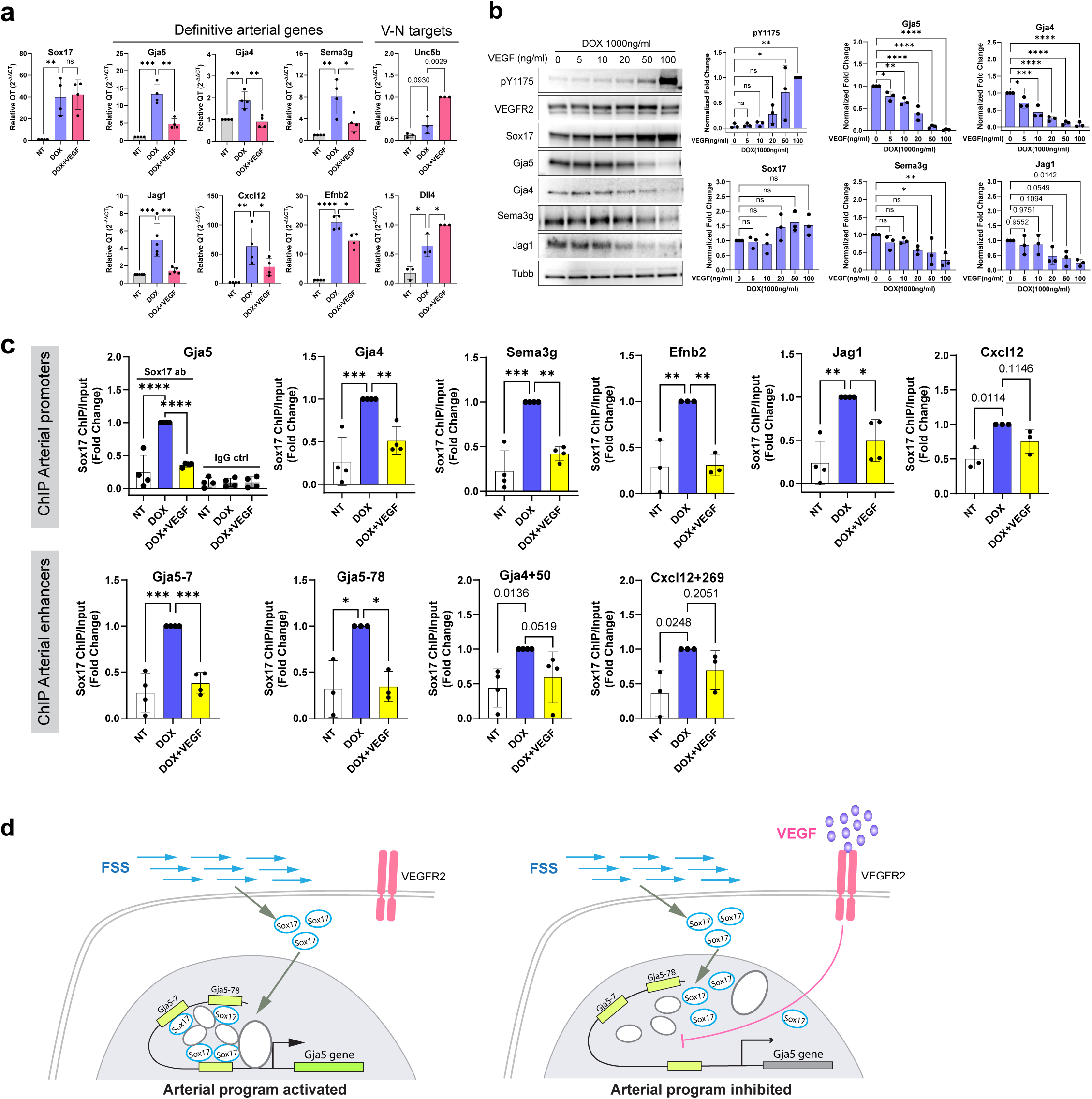
VEGF antagonizes Sox17 transcription activity on arterial promoters and enhancers. a-c. HUVECs harboring a doxycycline-inducible Sox17 expression system (pCW57.1-hSox17-myc-DDK) were used for these experiments for 48 hours. **a.** Quantitative RT-PCR analysis of RNA expression under experimental conditions of no treatment (NT), DOX (1000ng/ml) and DOX (1000ng/ml)+VEGF (50ng/ml). Definitive arterial genes (Gja5, Gja4, Sema3g, Jag1, Cxcl12 and Efnb2) and VEGF-Notch (V-N) targets (Unc5b and Dll4) were tested. **b.** A dose gradient of VEGF (0-100ng/ml) was used to test the impact on DOX-induced arterial differentiation. Cell lysates were analyzed by western blot. **c.** Chromatin immunoprecipitation (ChIP) results of Sox17 enrichment on the promoters and enhancers of the definitive arterial genes. The results are demonstrated by fold changes of percentage input relative to NT. **d.** Working model: VEGF signaling counteracts FSS-induced arterial specification by compromising Sox17-mediated transcription regulation. Gja4 is used as an example in this diagram. Left: in the absence of VEGF signaling, FSS-induced Sox17 is enriched at its binding motifs on arterial promoters and enhancers, forming an activating transcription complex to turn on arterial gene expression. Right: VEGF signaling compromises the enrichment of Sox17 on arterial promoters and enhancers, thereby turning off the expression of arterial gene. One-way ANOVA with multiple comparisons was used for statistics. A p value equal to or less than 0.05 was determined as statistically significant, in extents demonstrated by asterisks * (p≤0.05), ** (p≤0.01), *** (p≤0.001), **** (p≤0.0001). ns: not significant.

Since VEGF antagonizes Sox17-dependent genes without reducing Sox17 expression in HUVECs (Fig. 8a-b, Fig. S1), we examined Sox17’s transcriptional activity in arterial promoters and recently reported arterial enhancers^36^ using a chromatin immunoprecipitation (ChIP) assay to pull down Sox17-bound chromatin in the Dox-Sox17-HUVEC system. Cells were either subjected to no treatment (NT), Dox alone or a Dox+VEGF combination (Fig. 8c). As expected, Dox-induced Sox17 expression led to enrichment of Sox17 binding on arterial promoters and enhancers, including the proximal promoters of Gja5, Gja4, Sema3g, Jag1, Cxcl12 and Efnb2, and arterial enhancers Gja5-7, Gja5-78, Gja4+50, Cxcl12+269 (Fig. 8c), which was effectively inhibited by VEGF (Fig. 8c). VEGF thus counteracts the FSS-induced arterial program by compromising the transcription activity of Sox17 on arterial genes (Fig. 8d).

## Discussion

In summary, studies presented above show how angiogenesis and FSS interact to balance vascular growth and arterial patterning via the control of transcription factor Sox17 that functions as the key driver of capillary-to-arterial fate transition. FSS induces arterial fate specification in capillary endothelial cells, which is counteracted by VEGF. Therefore, the condition with high VEGF and low FSS favors angiogenesis, while the one with low VEGF and high FSS favors arterial patterning (Fig. 9a).

**Figure 9:**
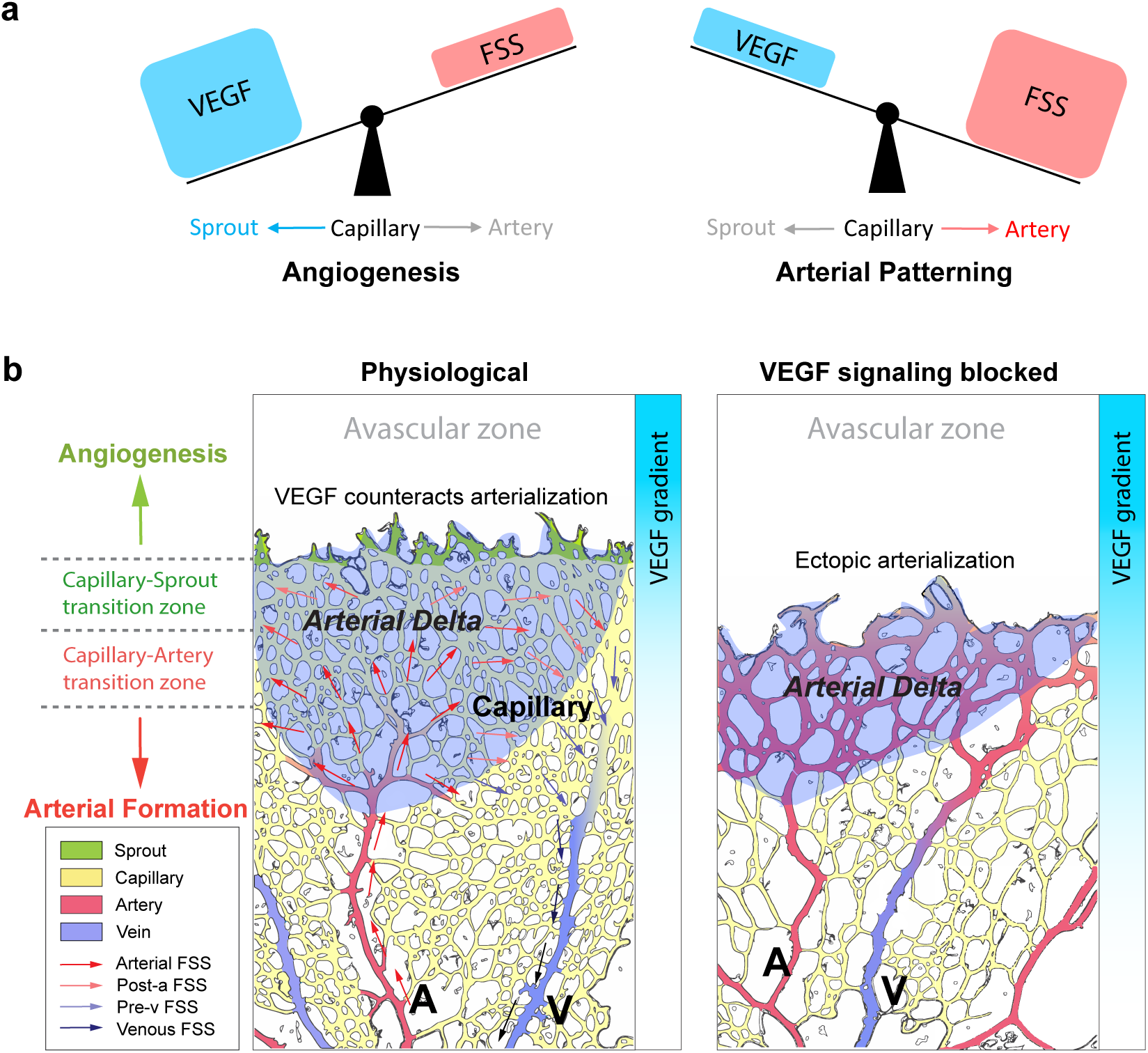
Angiogenic signaling is a physiological brake to counteract FSS-driven arterial patterning. **a.** VEGF and FSS counteract to control angiogenesis and arterial patterning from the capillary bed. The condition with high level of VEGF and low FSS favors angiogenesis and inhibits arterial formation, while arterial formation requires a decline of VEGF signaling as a permissive prerequisite to allow high arterial level of FSS to drive arterial specification. **b.** Artistic modeling of postnatal retinal vascular morphogenesis under physiological condition or blocked VEGFR2 signaling. Red arrows indicate the directionality of arterial (dark red) and post-arterial (light red) FSS. Blue arrows indicate the directionality of pre-venous (light blue) and venous (dark blue) FSS. The shaded artery-associated capillary area between the arterial tip and the vascular front is defined as the “*Arterial Delta*” in this study. Exposed to effective post-arterial FSS from the artery and high level of VEGF diffused from the juxtaposed avascular zone, the un-arterialized Arterial Delta region of capillary is where VEGF-FSS interaction takes place. Blocking VEGF signaling removes the physiological brake that counteracts the FSS-induced arterial program, leaving FSS as the sole player that ectopically arterializes the Arterial Delta and beyond.

These conclusions are supported by several lines of evidence. First, in vitro, FSS induces arterial gene expression in HMVBECs and HUVECs that is inhibited by VEGF in a dose-dependent manner. Second, developing retinal vasculature in vivo shows a clear spatiotemporal segregation of angiogenesis (early, on the periphery, in the area of high VEGF levels) versus arterial formation (late, central, in the areas of low VEGF levels). Suppression of VEGF/VEGFR2 signaling, by VEGF, VEGFR2, and PLCγ1 knockouts or with a VEGF neutralizing antibody (DC101), resulted in variably impaired angiogenesis but consistently preserved arterial formation with ectopic arterialization (Fig. 9b). VEGF is thus required for forming and expanding the capillary bed, as expected^37–39^, but FSS-driven capillary remodeling into arterial vasculature is independent of VEGFR2 signaling. Importantly, VEGF serves as a physiological brake to counteract FSS-driven arterialization, thus maintaining the capillary bed and preventing ectopic arterialization, while cessation of VEGF-VEGFR2 signaling is required for FSS-driven arterial specification in the central region (Fig. 9b).

This explains the lack of arterialization in the artery-associated capillary region (Arterial Delta) where the two environmental stimuli (VEGF and FSS) interact, which demonstrates a spatiotemporal segregation of vascular growth and patterning (Fig. 9b). Blocking VEGF signaling removes the physiological brake for FSS-driven arterial specification, leading to excessive arterialization and loss of a functional capillary bed in the Arterial Delta (Fig. 9b, right panel). This also explains why ectopic arterialization takes place in the peripheral but not central part of the capillary bed upon loss of VEGF signaling.

Our study identifies the key signaling mechanism for VEGF inhibition of arterial fate specification. The SoxF family transcription factors, Sox7, Sox17 and Sox18, are expressed by all ECs^34^. Among them, only Sox17 is enriched in arterial ECs and is implicated in arterial specification^17,34^. Our loss-of-function and gain-of-function data show that Sox17 is a critical mechanosensitive transcription activator that mediates the FSS-arterial program. Importantly, the transcription activity of Sox17 is sensitive to the inhibitory effect of VEGF signaling that counteracts the FSS-arterial program. This transcriptional regulation mechanism establishes a molecular basis for the new paradigm addressed above. Exactly how FSS mechanotransduction activates and VEGF signaling blocks Sox17 transcriptional activity are key questions for further studies.

Previous studies proposed that VEGF induced Dll4 expression, which then activates Notch signaling to induce arterial specification^10,12,13^. It is well established that VEGF-induced Notch signaling functions primarily as a negative regulator to stop VEGF signaling^8,18,20^ and arrest cell cycle progression^21^, thereby terminating angiogenesis and proliferation to control vascular density. This negative feedback mechanism in the endothelium, together with the cessation of VEGF production from the vascularized neural bed, spatially limits VEGF signaling to the dynamically moving vascular front and avoids persistent VEGF signaling after tip-to-stalk conversion. Although constitutively active Notch can induce arterial specification^17,21^, it is unlikely that transient VEGF signaling at the angiogenic front leads to persistent Notch signaling that subsequently induces arterial fate in the non-sprouting vasculature. Moreover, it has been proposed that the tip cells that competitively acquire high VEGF-Notch signaling are partially committed to arterial program as “pre-arterial” ECs at the vascular front^16,40^. Our data shows that VEGF signaling is not required for arterial formation and that flow can induce a full arterial program independent of VEGF signaling as the primary mechanism. However, this may not exclude the presence of partially committed arterial progenitors at the angiogenic front. In fact, our proposed new concept of VEGF’s inhibitory role counteracting arterial fate in this study could provide an explanation why these “pre-arterial” ECs, if present, don’t undergo full specification at the angiogenic front and the Arterial Delta region of capillaries where VEGF levels are high.

In this study, we analyzed arterial morphogenesis in a comprehensive set of VEGF LOF mice. While each intervention led to distinct phenotypes, in all cases arterial formation was maintained with ectopic arterialization. In particular, DC101 treatment led to extensive ectopic arterialization in the arterial delta region. This likely reflects incomplete blocking of VEGF-A-VEGFR2 signaling and thereby better preservation of the capillary bed. Also, DC101 does not block ligand-independent functions of VEGFR2 such as mechanosensing and other biological activities driven by its numerous co-receptors and partner proteins such as VEGFR1, integrins, Nrp1, and Sdc2 among others^41–49^. While *VEGFR2^iECKO^*potentially jeopardizes other essential non-tyrosine kinase signaling pathways of VEGFR2, DC101 treatment leaves these functions intact. Furthermore, severe vascular rarefaction in *Vegfa^iKO^*, *Plcg1^iECKO^* and *VEGFR2^iECKO^* mice likely impairs cardiac output, resulting in decreased FSS, and creates uneven blood flow in the regressing vessels, further weakening the magnitude of the FSS. By contrast, the preservation of a substantial capillary bed in the DC101-treated retinas provides a more permissive vasculature with effective FSS. These findings also illustrate the need of angiogenesis for formation of an adequate capillary bed to support effective arterial morphogenesis.

While the arterial vasculature in *Vegfa^iKO^*and *Plcg1^iECKO^* mice retinas show similar magnitudes of Sox17^high^-labeled arterial extension, there are distinct differences in the extent of smooth muscle coverage in *Vegfa^iKO^* compared to *Plcg1^iECKO^* mice. This difference likely reflects a more profound inhibition of VEGF-A-VEGFR2 signaling in the absence of VEGF (*Vegfa^iKO^*) than PLCγ1 deletion, leading to different preservation of the capillary bed for arterial patterning. As PLCγ1 knockout blocks the signaling from VEGFR2-Y1173, signaling from other tyrosine phosphorylation sites of VEGFR2, such as Y951, Y1054, Y1059, Y1214, is retained. Furthermore, the uncoupling of arterial specification from mural smooth muscle coverage in the *Vegfa^iKO^* retinas suggests defective pericyte physiology and EC-pericyte crosstalk, as a global loss of VEGF also affects pericytic VEGFR1 signaling essential for normal pericyte biology^28,29^.

In conclusion, this study elucidates the coordination of vascular growth and patterning by the dynamic interaction of VEGF and FSS to create properly organized vasculature. Transient VEGF stimulation is necessary for the formation of the capillary bed, which FSS then partially remodeled into the arterial vasculature. The key finding that this latter step requires cessation of VEGF signaling has important implications for therapeutic and engineering revascularization, a critical area for future work.

## Methods

### Animal studies

*Vegfr2^flox/flox^* mice (a gift kindly provided by Lena Claesson-Welsh, Uppsala University), *Plcg1^flox/flox^* mice (a gift kindly provided by Renren Wen, Wisconsin Blood Institute) and *Dll4^flox/flox^* mice (a gift kindly provided by Prof. Bin Zhou, Albert Einstein College of Medicine) were crossed with *Cdh5CreER^T2^*mice (a gift kindly provided by Ralf Adams). *CBF:H2B-Venus* mice, purchased from Jackson Laboratory, were crossed with *Vegfr2^flox/flox^; Cdh5CreER^T2^ mice and Plcg1^flox/flox^; Cdh5CreER^T2^ mice*, respectively*. Vegf2^flox/flox^; Cdh5CreER^T2^* mice were in *C57Bl/6J* background*. Plcg1^flox/flox^; Cdh5CreER^T2^ mice were backcrossed to C57Bl/6J* background for 10 generations. Gene deletion was induced by three consecutive dosages of 100µg Tamoxifen via daily peritoneal injection. Tamoxifen powder (Sigma, T5648) was suspended in corn oil (Sigma, C8267) and then dissolved by agitation at 37°C for 2 hours(hrs). Rat IgG or DC101 antibody was injected peritoneally at postnatal day 1 (400µg) and 3 (200µg). All aforementioned mouse experimental procedures complied to the protocols approved by the Yale University Institutional Animal Care and Use Committee (IACUC). Genotyping primers are listed in Table S4. The following study was performed at the UCL Institute of Ophthalmology following UK Home Office and institutional Animal Welfare and Ethical Review Body (AWERB) guidelines. Mice carrying the *Vegfa*^fl/fl^ alleles^38^ maintained on a mixed genetic background (C57BL6/J and 129/Sv) were crossed with *Vegfa*^fl/fl^ mice carrying a tamoxifen-inducible *Cre* transgene under the control of the ubiquitous chicken actin promoter (CAG-CreER^TM^)^50^. *Vegfa*^fl/fl^ littermate pups expressing or lacking CAG-CreER^TM^ received intraperitoneal injections of 50 µg tamoxifen on postnatal days (P) 2 and P4.

### Retinal dissection

Mouse neonates were euthanized at postnatal day 1(P1) to 6(P6). Eyes were enucleated and immediately fixed with 4% paraformaldehyde (PFA) for 8 minutes (min) at room temperature (RT). Then the eyes were transferred to PBS to terminate pre-dissection fixation. Dissected retinas were fixed with 4% paraformaldehyde for 30 min at RT with gentle rocking, followed by 2 washes in PBS at RT, 10 min each. Then retinas were dehydrated in methanol and stored at -20°C for subsequent procedure. Before immunostaining, retinas stored in methanol were warmed up at room temperature and rehydrated with PBS.

### Immunostaining and confocal imaging

Retinas were incubated in wash buffer (PBS with 0.1%Triton X100) with gentle rocking for 15 min and then in blocking buffer (wash buffer with 5% normal donkey serum (Jackson ImmunoResearch, 017-000-121)) for 1-2 hrs. Blocked retinas were incubated with primary antibodies diluted in blocking buffer overnight at 4°C with gentle rocking. On the next day, the retinas were washed in wash buffer for 3 times, 15 min each. Secondary antibodies were mixed with isolectin B4 in blocking buffer for staining at RT for 1.5 h with gentle rocking in dark. Staining procedure was finished after 3 washes in wash buffer, 15 min each. The retinas were transferred to PBS without detergent before flat mount to slides. Antibodies and experimental conditions were listed in Table S5. Image acquisition was conducted with Leica TCS SP8 Confocal Laser Scanning Microscope. Leica LAS-X, Image J (NIH) and IMARIS (Bitplane, USA) were used for image analysis.

### EdU labeling

For in vivo EdU incorporation, 50ug EdU (Sigma #900684) solution were peritoneally injected into mouse neonates at P5 4h before retinal dissection. Erg antibody (Abcam, # ab214341) was stained to visualize endothelial nuclei, followed by detection of EdU incorporation using EdU Cell Proliferation Kit (Invitrogen, #C10340).

For In vitro experiments, EdU was added to the media to achieve a final concentration of 5uM 2 hours before cell harvesting. Click-iT™ EdU Cell Proliferation Kit for Imaging (Invitrogen, C10340) was used to detect EdU incorporation with counterstaining of DAPI.

### Immunostaining of phospho-ERK1/2

Mouse neonates at P5 were placed on a warm pad at 37°C for 30min. The eyes were enucleated and immediately fixed in 4% PFA for 8 min, RT. Using ice-cold PFA will decrease fixation efficacy and must be avoided at this step. The retinas were then transferred to and dissected in ice-cold PBS. Post-dissection fixation in 4% PFA was done at 4°C with gentle rocking for 2 h. The fixed retinas were washed once with PBS for 5min, RT, and stored in ice-cold methanol at -20°C for 1 h. After warming up at RT for 15min, the retinas were rehydrated in PBS for 5min, followed by incubation in PBST (PBS with 0.25%Triton X100) for 30min and then blocking in PBST with 10% normal donkey serum (blocking buffer) for 1hr, RT. Phospho-p44/42 MAPK (Erk1/2) (Thr202/Tyr204) antibody (Cell signaling Technology, #9101) were diluted 1:100 in blocking buffer. From this step, the staining procedure followed the regular protocol described above.

### Vascular perfusion assay

Mouse neonates at P5 or P6 were placed on a warm pad at 37°C for subsequent procedure of vascular perfusion. IB4-Alexa488 was diluted in PBS and retro-orbitally injected to visualize patent (lumenized) vessels. Calculation of IB4 dilution (1:50) was based on estimated blood volume 5.5ml/100g body weight (The Jackson Laboratory). 1:50= (IB4 volume) : (blood volume + injection volume). Injected pups were placed on warm pad for 20min before dissection. Dissected retinas were counter-stained with IB4-Alexa 647 to visualize all vessels, lumenized and unlumenized.

### Endothelial cell culture and flow (shear stress) systems

Human microvascular blood endothelial cells (HMVBECs) (HDBECs, Promocell, #C-12211) were maintained in EGM2-MV full growth medium (Lonza). EBM2 basal medium with 5% FBS, here we term *permissive arterial differentiation medium (PADM5)*, was used for flow-induced arterial differentiation. Approximately 6 h prior to start of flow, EGM2-MV medium was removed, followed by a wash and a refill with PADM5. Human Umbilical Vein Endothelial Cells (HUVECs) (from Yale VBT) were maintained in M199 medium containing 20%FBS and endothelial cell growth supplements (ECGS, aliquots of Corning product were purchased from Yale VBT). For HUVECs, the maintenance medium was used for flow-induced arterial differentiation.

HMVBECs or HUVECs were seeded in IBIDI µ-slides (0.4 Luer ibiTreat) (IBIDI, #80176), with an initial seed density of 0.15x 10^6^ (HMVBECs) or 0.12x 10^6^ (HUVECs) cells per slide and were maintained for 48 h in the slide channel to reach a monolayer of homeostasis before flow experiment.

Flow system was built with a mini peristaltic pump (Cole Parmer MasterFlex Peristaltic Pump C/L drive P/N: 77122-22), an assembly of tubing systems with a syringe pulse dampener and a medium reservoir. The tubing system was assembled with a coiled inlet tubing (60cm) and a coiled outlet tubing (160cm), a total length of 220cm to further dampen pulsation. To avoid cell detachment, the end of the outlet tubing was fit into the pump head to passively draw the flow out of the IBIDI channel instead of pumping in. A flow rate of 12ml/min, which creates 12 dynes/cm^2^ shear stress according to the IBIDI manual, was accurately measured in each flow system prior to the start of flow experiment. Before connecting IBIDI µSlides, the flow systems were placed in cell culture incubator for 30min for equilibrium of temperature and CO_2_.

To harvest cells from IBIDI µSlides for RNA or protein analysis, 150µl lysis buffer was added into the channel, followed by a freezing-thawing cycle to facilitate cell lysing. To collect cell lysate from the channel, approximately 75µl of thawed lysate was removed in order to create room for liquid movement. Then the slide was gently tilted 10-15 times to make the rest cell lysate moving through the channel back and forth. After collecting the rest cell lysate, 50ul of new lysis buffer was added into the channel and then passed through repeatedly by gentle tilting to rinse off the residual cell lysate. A total of 200ul cell lysate was collected from each channel.

### Small interfering RNA (siRNA) experiments

FuGene SI transfection reagent (VWR, # MSPP-SI-1000) was used for siRNA transfection for lower toxicity. HUVECs or HMVBECs were seeded in maintenance medium to reach a density of 80-90% on the day of transfection. siRNA/FuGene/OptiMEM mix was added to EGM2-MV full growth medium for 6h, followed by a wash with PBS and replacement with fresh EGM2-MV full medium. On the next day, the siRNA-treated cells were trypsinized and seeded in IBIDI µSlides with maintenance medium. Flow experiments started at 72h after siPLCγ1 transfection. siRNA information is listed in Table S6.

### RNA isolation and RT-qPCR

Qiagen RNeasy plus Mini kit (#74134) was used for total RNA isolation, followed by reverse transcription to synthesize cDNA using iScript cDNA Synthesis Kit (Bio-Rad, #170–8891). SYBR Green Supermix (Bio-Rad, #1798880) was used for qPCR. Cycle threshold (Ct) values of gene expression were subtracted from average housekeeping gene GAPDH, and relative quantification of experimental groups versus control was calculated. QPCR primers are listed in Table S7.

### RNA sequencing and bioinformatic analysis

Total RNA samples were sent to Yale Center for Genomic Analysis (YCGA) for quality analysis, library preparation and sequencing. Partek Flow was used for bioinformatic analysis. Briefly, adapters in RNAseq reads were trimmed, and the trimmed reads were aligned to aligner algorithm STAR. Normalized gene counts were analyzed using DESeq2 to determine differential gene expression.

### Computational methods and cSTAR analysis

To analyze EC responses to FSS, VEGF and a combined FSS plus VEGF, the control condition is selected as no treatment, static condition (ST) and the log fold-changes of the differentially expressed genes with respect to ST were calculated. The SVM algorithm with a linear kernel from the scikit-learn python library was applied to build a maximum margin hyperplane that distinguishes four different EC states. The STVs are calculated as unit vectors, which are normal vectors to the three hyperplanes separating EC states. After building the STVs, cSTAR calculates quantitative indicators of cell phenotypic states termed DPDs. The DPD scores describe the changes in the cell phenotypic features that would occur when the data point crosses on the separating hyperplanes. The absolute value of the DPD score is determined by the Euclidean distances from the data point to the separating hyperplane, while the sign of the DPD score indicates if the direction of the data point movement to cross the hyperplane is parallel or antiparallel to the corresponding STV^24^. The DPD scores were normalized so that the DPD values in the training set (FSS, VEGF, and FSS-VEGF states) were -1, 0 or 1. All codes of building STVs and calculating the normalized DPD scores will be provided in a journal publication.

### Isolation of mouse retinal endothelial cell and scRNAseq

Retinas of control and *PLCγ1^iECKO^* pups at p6 were dissected in ice-cold PBS. Dissected retinas from 3 control and 3 *PLCγ1^iECKO^* pups, respectively, were pooled and subjected to magnetic-activated Cell sorting (MACS). Briefly, pooled retinal tissue was dissociated using Neural Tissue Dissociation kit (P) (Miltenyi Biotec, Cat#130-092-628) with gentleMACS Dissociator (Miltenyi Biotec, Cat#130-093-235). Isolated cells were suspended in PBS with 0.4% ultra-pure BSA as single cell suspension, which was subsequently incubated with CD31 MicroBeads (Miltenyi Biotec, Cat#130-097-418) for 30 min at 4°C with gentle rotation followed by passing through MS columns (Miltenyi Biotec, Cat#130-042-201) on an OctoMACS Separator (Miltenyi Biotec, Cat#130-042-109) to enrich endothelial cells.

Single cell suspensions were submitted to Yale Center for Genomic Analysis (YCGA) for library preparation and sequencing. scRNAseq libraries were created using Chromium GEM-X Single Cell 3′ Reagent Kits v4 (10x Genomics). Libraries were sequenced in NovaSeq HiSeq paired-end.

Reads alignment and UMI counting were conducted using the Cell Ranger software suite, with the mm10 genome assembly utilized for alignment^51^. Downstream analyses, including quality control (QC) filtering, batch correction, clustering, UMAP visualization, cell cycle analysis, and gene expression analysis, were performed in R using Seurat v5.1^52^. For quality control, cells with mitochondrial gene content exceeding 5% were excluded. Additionally, only cells with fewer than 100,000 total counts and between 200 and 9,000 detected features were retained for analysis. Counts normalization was executed using the Pearson residuals method implemented in the sctransform package^53^. Batch correction and data integration were carried out using the RPCA approach available in Seurat package^52^. TSCAN method was employed for pseudotime trajectory analysis^54^. For RNA Velocity Analysis, splicing-specific counts were generated using the velocyto.py package^55^. For downstream processing of splicing data, the velocyto.R and Seurat-Wrappers packages were utilized^55^. Vector field plotting was conducted using the veloviz package^56^ to visualize RNA velocity dynamics.

To access endothelial deletion of Plcg1, the brain tissue of control and *PLCγ1^iECKO^* pups were used to isolate ECs. Brain ECs were first enriched by MACS as described above, then CD31+/CD45-/PI- ECs were specifically sorted out by fluorescence activated cell sorting (FACS) for RNA isolation and RT-PCR. The FACS-related antibodies are summarized in Table S5. Deletion of Plcg1 in *PLCγ1^iECKO^*ECs is evidenced in Fig. S4, which demonstrates the deletion of LoxP-flanked exons2-4 and the presence of a truncated Plcg1 mRNA with the downstream exons (Fig. S4b). This explains the observation of no detectable decrease of Plcg1 mRNA in the *PLCγ1^iECKO^* ECs in scRNAseq analysis (Fig. S4c), as Chromium GEM-X Single Cell 3′ Reagent Kits v4 uses the poly(dT) primers that capture the polyA sequences of mRNA and generates cDNA libraries with downstream sequences at the 3’ end, similar to a recent scRNAseq study with the same issue to detect gene deletion^57^.

### Western blots

The following recipe (kindly provided by Prof. Jannifer Fang, Tulane University) was used to prepare 10ml lysis buffer for detecting Connexins (Gja4 and Gja5) and also workable for other proteins. A tablet of protease inhibitor cocktail (Roche) and a tablet of PhosSTOP were dissolved in 4.75ml water. Then the following reagents were sequentially added and then mixed: 1.25ml 1M Tris, 2ml 20%SDS, 2ml Glycerol, 1ml beta-mercaptoethanol and a tip amount of blumophenol blue. At the end of flow experiment, IBIDI µSlides were detached from the flow system and placed on ice. To harvest protein lysate from IBIDI µSlide, medium was vacuum removed from the channel and replaced with 150ul lysis buffer. To enhance protein lysing, the IBIDI µSlide with lysis buffer was subject to a freezing-thawing cycle. Subsequently, with the removal and collection of 50% lysis buffer to create room for movement in the channel, the slide was repeatedly tilted several times to create shear force to enhance protein extraction before collecting the rest protein lysate. An additional 50ul of new lysis buffer was pipetted into the channel to rinse and collect any residual protein lysate, making a total volume of 200ul. Protein lysate was collected in a centrifuge tube and sonicated for 1 min at 15% power, followed by heating at 75°C for 30 min to denature protein. Antibodies used for western blot were listed in Table S8.

### Doxycycline-induced Sox17 expression and induction of arterial differentiation

Doxycycline-inducible Sox17 expression lentiviral construct pCW57.1-hSox17-DDK-Myc was created in the Simons lab and can be shared upon reasonable request. Maintained in the aforementioned M199-based media (M199 supplemented with 20%FBS and ECGS), HUVECs at passage 2-3 were used for Lentiviral transduction, during which EGM2-MV media was temporarily used to replace M199-based media to enhance transduction efficiency and cell survival. Transduced HUVECs were selected by 0.8ug/ml puromycin in EGM2-MV media for 48 hours, followed by a 48-hour window of recovery. Puromycin-selected, transduced HUVECs were either frozen for future experiments or seeded in M199-based media for subsequent arterial differentiation experiments.

To induce arterial differentiation, 1.9-2.0 x 10^6^ HUVECs harboring pCW57.1-hSox17 were seeded in M199-based media in each 10-cm dish on day1, which ensures the formation of a confluent endothelial monolayer for the start of doxycycline treatment on the next day (day2). Fresh media with doxycycline was replaced once on day3. The cells were harvested at the end of a 48-hour treatment on day4.

To ensure consistent and reproducible results, HUVECs must be regularly maintained in M199-based medium instead of EGM2, with final passage no more than 6 at the end of arterial differentiation.

### Chromatin Immunoprecipitation (ChIP)

ChIP assays were conducted using SimpleChIP Plus Enzymatic Chromatin IP kit (Cell Signaling Tech #9005) following the official protocol. Briefly, HUVECs in a 10-cm dish were fixed with ultra-pure formadehyde (methanol free) (Polysciences, 18814-10) to create protein-chromatin crosslink. Immunoprecipitation was performed with antibody targeting Sox17 (CST#81778) and control rabbit IgG (CST#2729). Pull-downed chromatin was eluted, purified and assayed by QPCR.

### CRISPR-cas9-mediated gene deletion

Oligos of designed gRNA sequences were cloned into lentiCRISPRv2 (Addgene #52961) following previously published protocol^58^. The following gRNA sequences were used, including a non-targeting gRNA (GTATTACTGATATTGGTGGG), a pair of gRNAs targeting human Sox17 (ACGGGTAGCCGTCGAGCGG and GGCACCTACAGCTACGCGC)^59^, and a gRNA targeting human Sox18 (GTGGAAGGAGCTGAACGCGG). HUVECs transduced with lentiviral-delivered sgRNAs were selected with 0.8ug/ml puromycin for 48hrs and maintained in EGM2-MV medium for 7 days to ensure sufficient gene recombination. This was followed by a second cycle of selection with puromycin for 48hrs and recovery for 3 days before seeding the cells to the IBIDI µ slides for flow experiments.

### Statistics

Data was plotted and analyzed using GraphPad Prism10. Student’s t test was used for comparison of two groups, with the assumption of normal distribution and equal variance between groups. One-way or two-way ANOVA with multiple comparisons was used for grouped samples on one or two variables. A p value equal to or less than 0.05 was determined as statistically significant.

## Author contributions

DC, MS and MAS conceptualized and designed the project. DC conducted and analyzed most of the experiments. MS and MAS supervised the project. DC and MS wrote the manuscript with edits from MAS, CR and KAM. OSR, DC, AT, BNK analyzed the scRNAseq data. DC analyzed the RNAseq data. OSR and BNK conducted the cSTAR analysis based on the RNAseq data. DJ and BGC contributed to the shear stress experiments and cloning the CRISPR-Cas9 system for Sox17 deletion and provided technical assistance for DC to create the lentiviral construct of Sox17 expression. DC and RC designed and conducted the Sox17 ChIP experiments with KAM’s supervision. MR and EI conducted the experiments of the VEGF knockout mice with CR’s supervision.

## Acknowledgements

We thank Prof. Lena Claesson-Welsh, Prof. Ralf Adams, Prof. Renren Wen and Genentech for providing the *VEGFR2^flox/flox^*, the *Cdh5CreER^T2^,* the *PLCγ1^flox/flox^* and the *Vegfa ^flox/flox^* mice, respectively. We thank Dr. Hyojin Park and Dr. Eunate Gallardo for essential technical support in sample preparation for single cell RNA sequencing. Sequencing service by Yale Center for Genomic Analysis reported in this publication was supported by the National Institute of General Medical Sciences of the National Institutes of Health under Award Number 1S10OD030363-01A1. We thank Drs. Prajwal Boddu, Emma Ristori and Federico Corti for technical advice on cloning. We thank Rolando Milian, Yale Medical Library, for bioinformatic training and assistance of RNAseq data analysis. We thank Tuo Guo and Guilherme Dantas for technical assistance in immunostaining, confocal imaging and cloning. This work was supported by NIH grants R01HL169520-01 (MS), P01HL107205-06A1 (MS and MAS), NIH/NCI grant R01CA244660 (BNK), EU grant no. 101136926 MULTIR (BNK), Science Foundation Ireland 22/PATH-S/10875 (OSR) and Open Philanthropy (MS), British Heart Foundation PG.23.11342 (CR and MR) and PG.24.11119 (CR).

