## Supplemental Figures and Tables for "Angiogenic Signaling Counteracts Shear Stress-driven Arterial Patterning"

### Supplemental Fig.1 VEGF-FSS crosstalk regulates arterial specification in HUVECs.

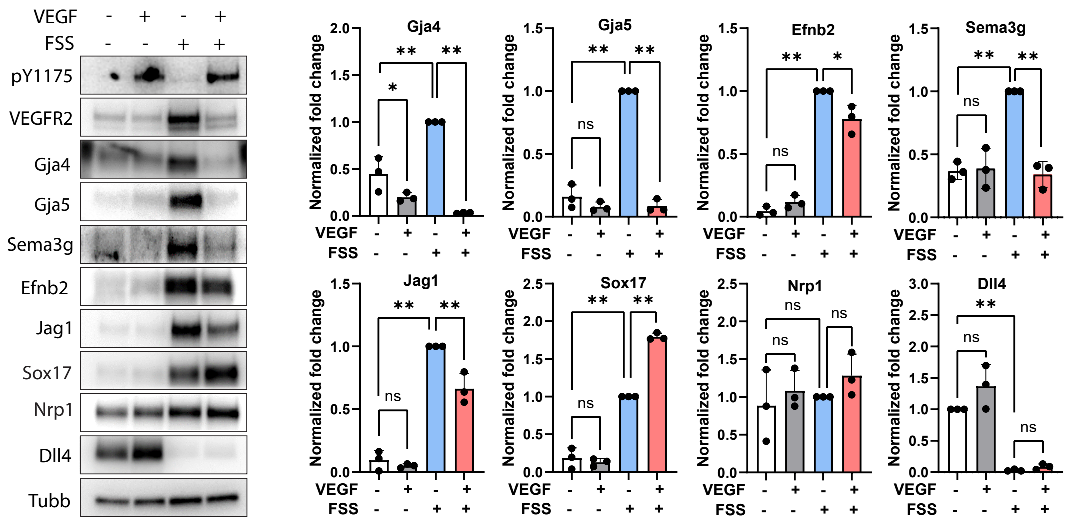

### Supplemental Fig.2 Endothelial deletion of PLCγ1 in P6 mouse neonates.

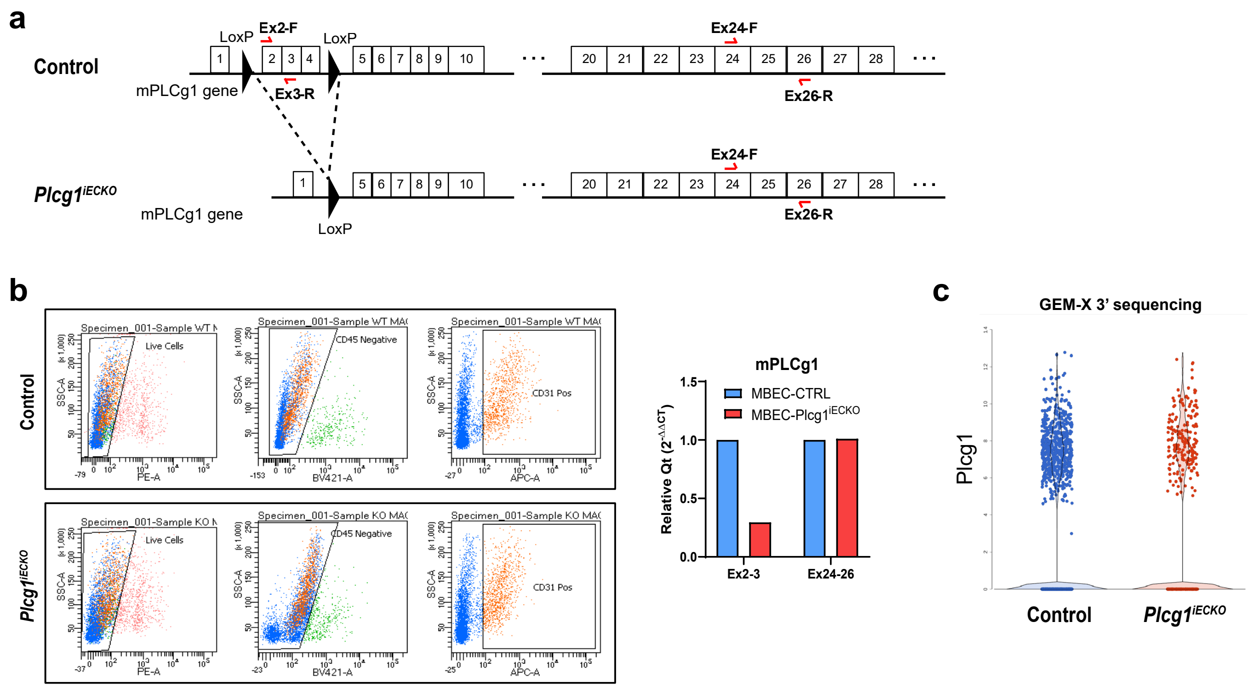

### Supplemental Fig. 3 Clustering of endothelial cells in a single-cell transcriptome.

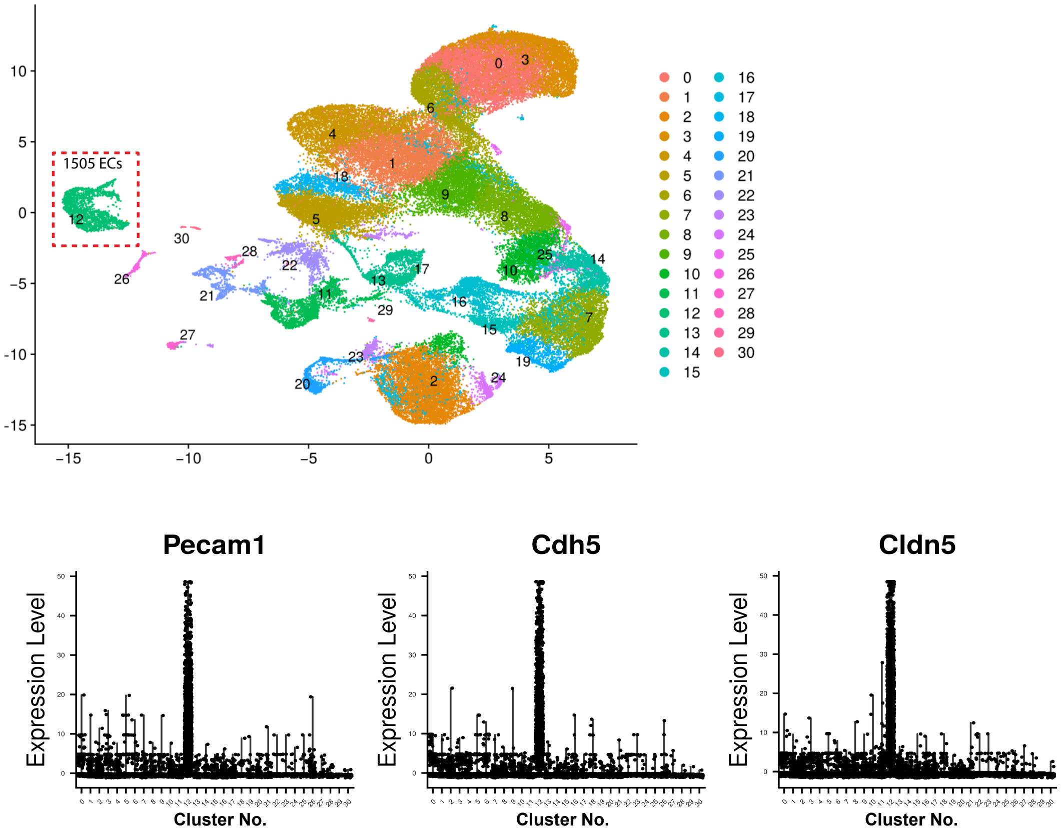

**Supplemental Fig. 4 Mapping arteriovenous zonation in a single cell transcriptome.**

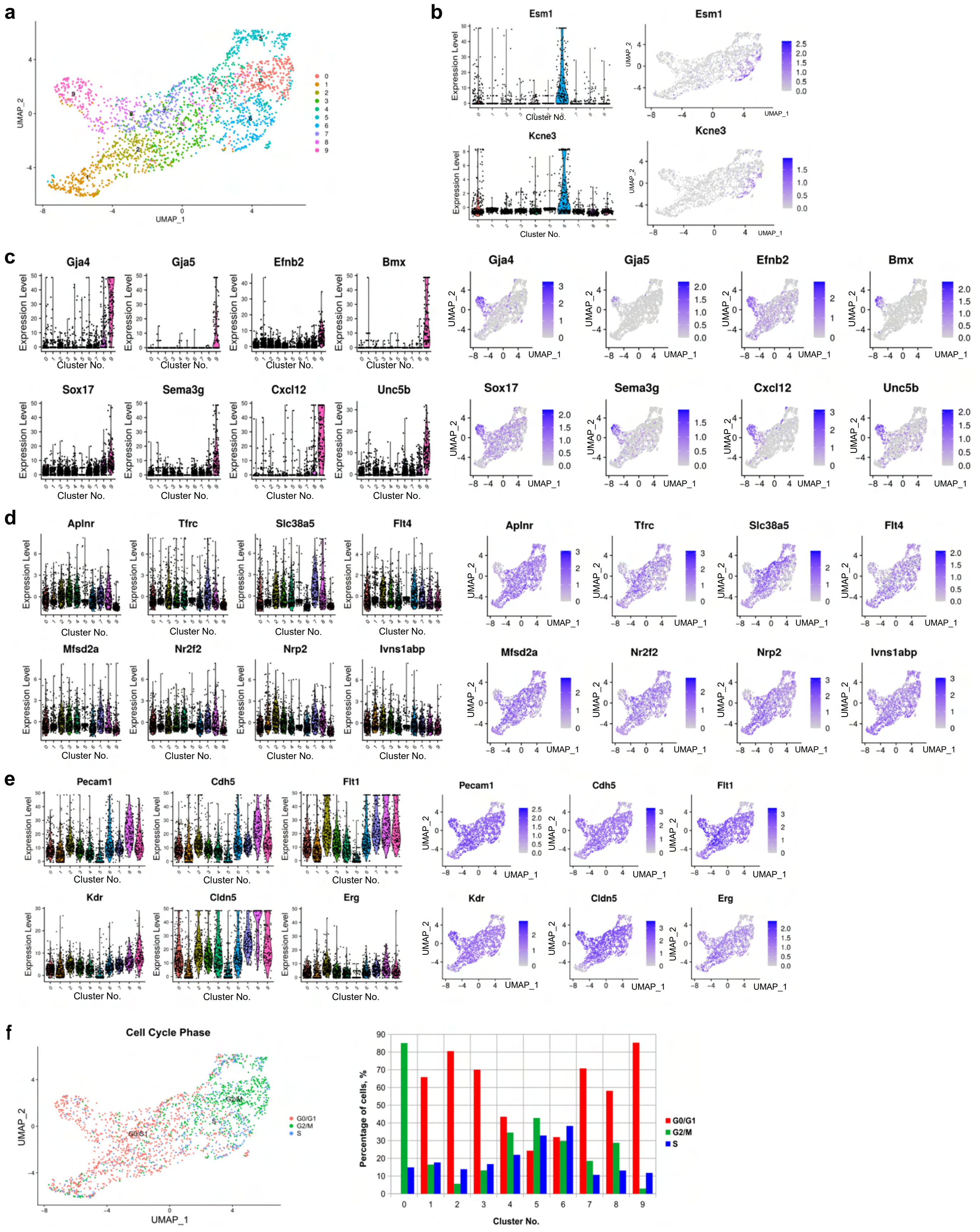

**Supplemental Fig.5 VEGF does not significantly alter FSS-induced cell cycle arrest.**

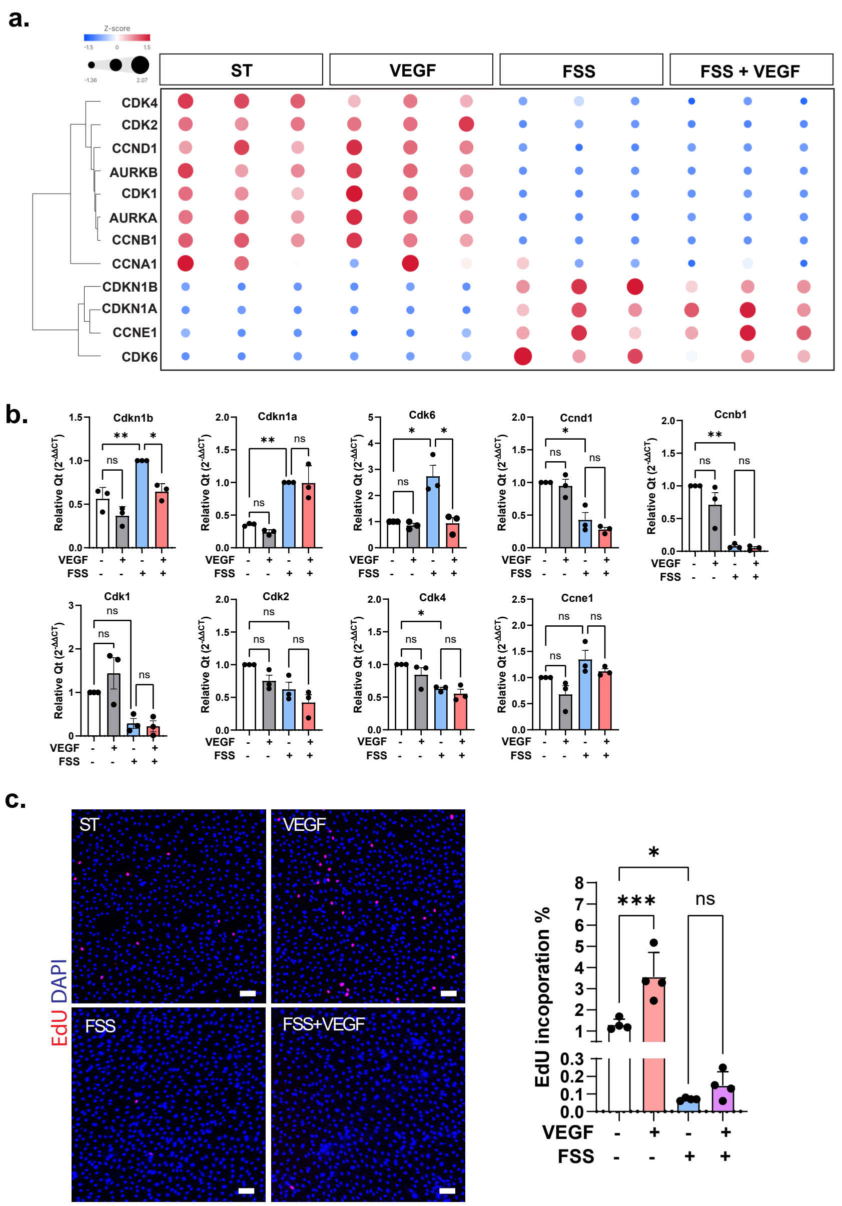

**Supplemental Fig.6 VEGF-A does not use Notch to counteract the FSS-induced arterial program.**

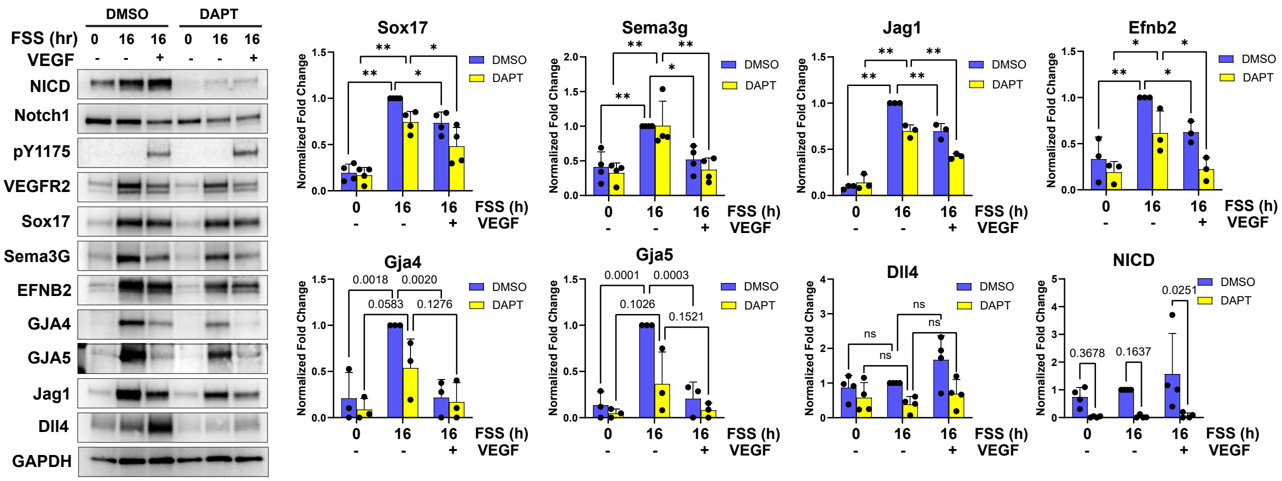

**Supplemental Figure Legend**

**Fig.S1. VEGF-FSS interaction regulates arterial specification in HUVECs.**
HUVECs were subjected to the conditions of static (VEGF-/FSS-), ST with 20ng/ml VEGF (VEGF+/FSS-), 12dyn/cm$^2$ FSS (FSS+/VEGF-) and FSS+VEGF (FSS+/VEGF+). Cell lyasates were analyzed by western blot. While most definitive arterial genes responded to VEGF and VEGF+FSS the same manner as HMVBECs (Fig.1), FSS induction of Sox17 in HUVECs is further promoted by VEGF. One-way ANOVA with multiple comparisons was used for statistics. A p value equal to or less than 0.05 was determined as statistically significant, in extents demonstrated by asterisks * ($p \leqslant 0.05$), ** ($p \leqslant 0.01$). ns: not significant.

**Fig.S2. Endothelial deletion of PLCγ1 in P6 mouse neonates.**
**a.** A demonstration of the exons in mouse PLCg1 gene in control and *Plcg1$^{iECKO}$*. Two loxP sites flank exons 2-4, which are excised in *Plcg1$^{iECKO}$*. Two sets of PCR primers were designed to detect Exons2-3 (Ex2-3) or downstream Exons 24-26 (Ex24-26). **b.** Pool brains from control and *PLCγ1$^{iECKO}$* pups at P6 (tamoxifen induction at P1-3) were first enriched by magnetic activated cell sorting (MACS) and further sorted by Fluorescence-activated cell sorting (FACS) in flow cytometry sorting out the CD31+ (APC)/CD45- (BV421)/PI- (PE) ECs. Plcg1 mRNA was detected by primer sets for Ex2-3 or Ex24-26. **c.** Distribution of retinal ECs with Plcg1 RNA expression in GEM-X 3' scRNAseq.

**Fig.S3. Clustering of endothelial cells in a single-cell transcriptome.**
30 clusters of cells from control and *Plcg1$^{iECKO}$* retinas were identified and demonstrated in UMAP. Endothelial cells in cluster 12 were identified by high expression levels of Pecam1, Cdh5 and Cldn5.

**Fig.S4. Mapping arteriovenous zonation in a single cell transcriptome.**
**a.** Identification of clusters 0-9 in all combined control and *Plcg1$^{iECKO}$* endothelial cells. **b.** Cluster 6 enriched with Esm1 and Kcne3 is identified as sprouting ECs. **c-d.** The cells in Cluster 9 expressing high level of arterial markers are arterial ECs. Clusters 0, 2, 3, 4, 6, 7 are identified as capillary/venous clusters. The cells in Cluster 8 are arterial predecessors with both arterial and capillary profiles. The ECs in Cluster 1 have weak capillary identity. **e.** The ECs in Cluster5 express low levels of endothelial markers, likely losing endothelial identity. **f.** Mapping cell cycle. Cluster 0 identified as proliferative capillary ECs mostly in G2/M phase.

**Fig.S5 VEGF does not significantly alter FSS-induced cell cycle arrest.**
**a.** RNAseq differential gene expression clustering analysis of cell cycle regulators. HMVBECs were cultured in the conditions of static (ST), VEGF (20ng/ml), FSS (12dyn/cm$^2$) and FSS+VEGF for 16 hours. Gene expression matrix is demonstrated by both color intensity and size. **b.** Validation of the arterial and capillary genes by quantitative RT-PCR. **c.** EdU incorporation assay. 5uM EdU was added to the media 2 hours before cell harvesting.

**Fig.S6. VEGF does not use Notch to counteract the FSS-induced arterial program.**
Pretreated with DMSO or 5uM DAPT for 3h, HMVBECs were harvested (FSS 0 hr) or subjected to FSS with DMSO or DAPT for 16 hr. Samples were analyzed by western blot. One-way ANOVA with multiple comparisons was used for statistics. A p value equal to or less than 0.05 was determined as statistically significant, in extents demonstrated by asterisks * ($p \leqslant 0.05$), ** ($p \leqslant 0.01$). ns: not significant.

**Table S1.** **Endothelial markers for mapping arteriovenous zonation.**

**Table S2. A matrix of the DPD scores.**

**Table S3. Predicted transcription factor binding motifs of Sox17 and RBPJ in arterial and capillary genes (TRANSFACT).**

**Table S4. Genotyping primers**

**Table S5. Antibodies for immunostaining.**

**Table S6. siRNA information.**

**Table S7. QPCR primers**

**Table S8. Antibodies for western blot.**

**Table S9. QPCR primers for ChIP.**

**Table S1. Endothelial markers for mapping arteriovenous zonation.**

| **Definitive Arterial genes** | **Unspecific arterial genes** (enriched in arteries and other EC subpopulations) | **Capillary genes** | **Sprouting markers** | **Venous markers** |
|---|---|---|---|---|
| Gja4<br>Gja5<br>Sema3g<br>Jag1<br>Sox17<br>Efnb2<br>Bmx<br>Gkn3<br>Cxcl12 | Unc5bB (A, S)<br>Nrp1 (A, C)<br>Dll4 (A, S) | Mfsd2a(C,V)<br>Flt4 (C,V, S)<br>Aplnr (C,V)<br>Tfrc (C,V)<br>Ivns1abp (C) | Esm1<br>Kcne3 | Nr2f2<br>Slc38a5<br>Nrp2 |

Abbreviations: A: artery. S: sprout. C: capillary. V: vein

**Table S2. A matrix of the DPD scores.**

| | **$DPD_{FSS}$** | **$DPD_{VEGF}$** | **$DPD_{SSVA}$** |
|---|---|---|---|
| **FSS** | 1.000 | -0.021 | 1.000 |
| **FSS-VEGFA** | 1.000 | -0.169 | -0.994 |
| **VEGF** | -0.948 | 1.000 | -0.929 |
| **STAT** | -1.000 | -1.000 | -1.000 |

**Table S3. Predicted transcription factor binding motifs of Sox17 and RBPJ in arterial and capillary genes (TRANSFACT).**

| Classification | Gene | Sox17 motif | RBPJ motif |
|---|---|---|---|
| Arterial | Sox17 | + | - |
| | GJA4 | + | - |
| | GJA5 | + | + |
| | EFNB2 | + | + |
| | SEMA3G | + | + |
| | NRP1 | + | + |
| Notch | HEY1 | + | + |
| | JAG1 | + | + |
| | DLL4 | + | + |
| | NOTCH1 | + | + |
| Capillary | MFSD2A | - | + |
| | FLT4 | - | + |
| | IVNS1ABP | - | + |
| | SLC16A11 | - | - |
| | TFRC | - | - |

#### Table S4. Genotyping primers

| Genotype | Forward primer (5'-3') | Reversed primer (5'-3') | Annealing Temp (°C) | Product Size (bp) |
|---|---|---|---|---|
| Cdh5creERT2 | CAGATCAGCTCCTCCACGAA | 1) TGGTGGGCAGGTAGCATGTT<br>2) CATTGCTGTCACTTGGTCGT | 55 | 356 (Cdh5cre)<br>144 (wt) |
| VEGFR2 flox | CCACAGAACAACTCAGGGCTA | GGGAGCAAAGTCTCTGGAAA | Touchdown (JAX 018977) | 230 (flox)<br>179 (wt) |
| PLCγ1 flox | TGTGTCAGTTTCATAGCCTGAA GAACGAG | TAAGAAAGCAGGCTGAGTAAGCC3 | 67 | 220 |
| PLCγ1 WT | ATCGTAACCGAGAGGATC | AGCAAGCCAGTAAGCCAGCACCC CTCCATG | 62 | 540 |
| Dll4 flox | GAGTCTGTCTGACTGACAGG | CTCGGTAGGTAATCCAGGTG | 58 | ~250 (flox)<br>~200 (wt) |
| Vegfa flox | CCTGGCCCTCAAGTACACCTT | TCCGTACGACGCATTTCTAG | | |
| Cag-creER$^{TM}$ | GCTAACCATGTTCATGCCTTC | AGGCAAATTTTGGTGTACGG | | |

#### Table S5. Antibodies for immunostaining and FACS.

| Antibody | Vendor | Catalogue # | Dilution |
|---|---|---|---|
| Emcn | R&D | AF4666 | 1:400 |
| Mfsd2a | Cell Signaling Tech | 80302 | 1:100 |
| Phospho-ERK1/2 | Cell Signaling Tech | 4370 | 1:100 |
| SMA | Sigma | C6198 | 1:500 |
| Sox17 | R&D | AF1924 | 1:100 |
| TFRC | R&D | AF2474 | 1:100 |
| VEGFR2 | BD Pharmingen | 555307 | 1:100 |
| VEGFR3 | R&D | AF743 | 1:100 |
| V450 Rat Anti-Mouse CD45 | BD Horizon | 560501 | 1:200 |
| APC Rat Anti-Mouse CD31 | BD Pharmingen | 551262 | 1:200 |
| APC Rat IgG2a κ Isotype Control | BD Pharmingen | 553932 | 1:200 |
| V450 Rat IgG2a, κ Isotype Control | BD Horizon | 560377 | 1:200 |

#### Table S6. siRNA information.

| siRNA | Vendor | Catalogue # |
|---|---|---|
| ON-Target plus Non-targeting Pool | Dharmacon | D-001810-10-05 |
| AllStars Negative Control | Qiagen | SI03650318 |
| siVEGFR2 | Qiagen | SI000605535 |
| siPLCγ1 | Qiagen | SI00041181 |

#### Table S7. QPCR primers

| Gene | Species | Forward primer | Reversed primer | Product size (bp) |
|---|---|---|---|---|
| VEGFR2 | Human | Qiagen, QT00069818 | | |
| PLCγ1 | Human | GGAAGACCTCACGGGACTTTG | GCGTTTTCAGGCGAAATTCCA | 108 |
| Sox17 | Human | CCAAAGCGGAGTCTCGCAT | GCCTAGCATCTTGCTTAGCTC | 125 |
| Efnb2 | Human | TTCTAGCACCGATGGCAACAG | CCCTGCGAATAAGGCCACT | 76 |
| Gja4 | Human | TGCAAGAGTGTGCTAGAGGC | ACAAAGCAGTCCACGAGGTAG | 119 |
| Gja5 | Human | CTGGCTCTTACGAGTACCCG | TGCACACATAGGTGTTGAGCA | 108 |
| Nrp1 | Human | ACCCAAGTGAAAAATGCGAATG | CCTCCAAATCGAAGTGAGGGTT | 90 |
| Sema3g | Human | Qiagen, 330001 PPH09633A | | |
| Sema3g | Human | CTGAGGAAGTGGTTCTGGAGGA | GCCGTAAGTCTCACATTGGTGC | 149 |
| Jag1 | Human | GTCCATGCAGAACGTGAACG | GCGGGACTGATACTCCTTGA | 136 |
| Cxcl12 | Human | ATTCTCAACACTCCAAACTGTGC | ACTTTAGCTTCGGGTCAATGC | 88 |
| Hey1 | Human | GTTCGGCTCTAGGTTCCATGT | CGTCGGCGCTTCTCAATTATTC | 88 |
| Notch1 | Human | GAGGCGTGGCAGACTATGC | CTTGTACTCCGTCAGCGTGA | 140 |
| Gapdh | Human | TGCACCACCAACTGCTTAGC | GGCATGGACTGTGGTCATGAG | 87 |
| S18 rRNA | Human | TAACGAACGAGACTCTGGCAT | CGGACATCTAAGGGCATCACAG | |
| PLCγ1$^{Ex2-3}$ | Mouse | GAGACGCGCCAGATCACAT | GCAGTGTGATTGATCTGGCC | 153 |
| PLCγ1$^{Ex24-26}$ | Mouse | ATCCAGCAGTCCTAGAGCCTG | GGATGGCGATCTGACAAGC | 105 |
| Gapdh | Mouse | AGGTCGGTGTGAACGGATTTG | TGTAGACCATGTAGTTGAGGTCA | 123 |

**Table S8. Antibodies for western blot.**

| Antibody | Vendor | Catalogue # | Dilution |
|---|---|---|---|
| Cx37 | ABclonal | A2529 | 1:1000 |
| Cx37 | Alpha Diagnostic Int. | CX37A11-A | 1:500 |
| Cx40 | ABclonal | A7231 | 1:1000 |
| Dll4 | Cell Signaling Tech | 2589 | 1:1000 |
| Efnb2 | R&D | AF496 | 1:500, fresh prep. |
| Emcn | R&D | AF7206 | 1:500 |
| GAPDH | Cell Signaling Tech | 5174 | 1:1000 |
| JAG1 | Cell Signaling Tech | 70109 | 1:2000 |
| Mfsd2a | Cell Signaling Tech | 80302 | 1:1000 |
| NICD | Cell Signaling Tech | 4147 | 1:1000 |
| NOTCH1 | Cell Signaling Tech | 3608 | 1:1000 |
| Nrp1 | Cell Signaling Tech | 3725 | 1:1000 |
| PLCγ1 | Cell Signaling Tech | 5690 | 1:1000 |
| Sema3g | Novus Bio | NBP1-83881 | 1:1000, fresh prep. |
| Sox17 | Cell Signaling Tech | 81778 | 1:1000 |
| Sox18 | R&D | AF5077 | 1:100 |
| Unc5b | Cell Signaling Tech | 13851 | 1:100 |
| VEGFR2 | Cell Signaling Tech | 9698 | 1:1000 |
| VEGFR2-pY1175 | Cell Signaling Tech | 2478 | 1:1000 |
| VEGFR3 | Cell Signaling Tech | 2638 | 1:1000 |

**Table S9. QPCR primers for ChIP.**

| Gene or enhancer | Species | Forward primer | Reversed primer | Product size (bp) |
|---|---|---|---|---|
| Gja5 | Human | GCAACCAACGAAAAGTTCGC | CCGTTAAAGTGTGGGAGCCA | 108 |
| Gja4 | Human | CCAGCCACCATTTAGAGGCA | GAGCAGGCGTCTCTATCCAC | 184 |
| Jag1 | Human | AGCATGCACGACTGGAAAAC | GGAGCGTCTCAAAGAAGCGA | 118 |
| Sema3g | Human | CAGGCTTTTGTGCCACACTG | CTCTGATCCCTGACCTCCCT | 156 |
| Efnb2 | Human | AAAGTCTCACATCCAAAGCGT | TCTTCGAAGTGAGTACTGCTGA | 213 |
| Cxcl12 | Human | TCATTCAGTTCCCGCCATCG | ACCTTAGGCGTAAAGTGGGG | 71 |
| Gja4+50 (hg38/chr1:34,842,968-34,843,574)[34] | Human | AAAAGGCCTGATAGCCGGTG | GACCAGCCAGGAATTCACCC | 281 |
| Gja5-7 (hg38/chr1:147,781,009-147,781,500) [34] | Human | GGCAGAGACAGCACAGAAGT | TAGGCATCCTCCCACTTCCA | 398 |
| Gja5-78 (hg38/chr1:147,877,925-147,878,475) [34] | Human | CCCAAGGGAGCAGTGCATAA | GCACAGTGCAAAGATGCCAA | 286 |
| Cxcl12+269 (hg38/chr10:44,043,453-44,044,439) [34] | Human | AGGATCTGCTTTGCAGAGGG | GGGCCAAACTAAAAGCCACG | 85 |
